# Deconstructing CTL-mediated autoimmunity through weak TCR-cross-reactivity towards highly abundant self-antigen

**DOI:** 10.1101/2024.08.17.608371

**Authors:** Angelika Plach, Vanessa Mühlgrabner, Aleksandra Rodak, René Platzer, Iago Doel Perez, Paul Fellinger, Timo Peters, Yvonne Winhofer, Christoph Madritsch, Lukas Schrangl, Hannes Stockinger, Loïc Dupré, Dirk H. Busch, Kilian Schober, Janett Göhring, Johannes B. Huppa

## Abstract

T-cell antigen receptors (TCRs) exhibit inherent cross-reactivity which broadens the spectrum of epitopes that are recognizable by a finite TCR-repertoire but also carries the risk of autoimmunity. However, TCRs support also a high level of antigen specificity as they allow T-cells to discriminate single antigenic peptide/MHC complexes (pMHCs) against millions of structurally related self-pMHCs, in some cases based on the absence or presence of a single methyl-group. How TCRs manage to convey such seemingly contrary properties and why some T-cells become over time autoreactive despite negative thymic selection, has remained elusive. Here, we devised a non-invasive molecular live cell imaging platform to investigate the biophysical parameters governing stimulatory TCR:pMHC interactions in settings of autoreactivity and anti-viral responses - two extremes in T-cell antigen recognition. We show that CMV-specific CD8+ RA14-T-cells respond effectively to even a single HLA-A2/CMV (A2/CMV) antigen, with synaptic TCR:pMHC lifetimes lasting seconds. In contrast, cross-reactivity of type 1 diabetes (T1D)-associated CD8+ 1E6 T-cells towards HLA-A2/preproinsulin (A2/PPI) self-epitopes involved ten-fold less stable synaptic TCR interactions resulting in severely attenuated ZAP70 recruitment and downstream signaling. Compared to A2/CMV-engaged RA14 T-cells, 1E6-T-cells required for activation 4000 or more A2/PPI and at least 100-times as many simultaneously pMHC-engaged TCRs. In support of antigen discrimination, CD8 co-engagement of MHC class I (MHCI) strengthened both settings of TCR:pMHC interactions equally but was essential only for sensitized virus detection but not autorecognition (1000-versus 5-fold enhancement). We conclude that the binding dynamics of TCRs and CD8 with pMHC shape the boundaries of central tolerance in the physiological context of the phenomenal yet also differential T-cell antigen detection capacity, TCR-cross-reactivity and self-antigen abundance. Gained insights are integral to a molecular and quantitative understanding of CD8+ T-cell mediated autoimmunity and protective immunity against infections and cancer.

**ONE SENTENCE SUMMARY:** With the use of newly devised molecular live-cell imaging modalities we measured with unprecedented precision T-cell antigen recognition dynamics in human T-cells in settings of anti-viral immunity and autoimmunity-causing cross-reactivity. These two extremes within the spectrum of T-cell antigen detection differed substantially with regard to synaptic TCR: antigen-engagement, the level of sensitization through the CD8-coreceptor and the overall efficiency of ensuing downstream signaling. Our results demarcate limits of central tolerance and protective immunity and set quantitative boundaries on the occurrence of autoimmunity with direct implications for T-cell-based designs of immunotherapies.

## INTRODUCTION

The ability to distinguish self-from non-self-antigens and to mount an efficient immune response while suppressing autoreactivity is a key characteristic of T-cell-based immunity ^1–3^. T-lymphocytes can sense the presence of even single stimulatory-pMHC molecules which are typically vastly outnumbered by structurally related yet non-stimulatory endogenous pMHCs on the antigen presenting cell (APC) surface ^4–7^. They meet this task with the use of genetically rearranged clonotypic T-cell antigen receptors (TCRs), which bind to antigenic pMHCs within the immunological synapse, the area of contact formed with an antigen presenting cell (APC). Both the clonotypic identity of the TCR as well as the primary structure of the MHC-presented peptide define the TCR:pMHC binding strength and the unique ability of T-cells to translate differences as subtle as the presence or absence of a single methyl group within the presented peptide (e.g. threonine versus serine residue) into diverging downstream signaling ^8,9^.

Due to the phenomenal processivity of their underlying cellular signaling machinery, most TCRs exhibit significant cross-reactivity towards unrelated peptides which raises invariably the risk of T-cell-mediated autoimmunity. As a safeguard, central tolerance mechanisms eliminate self-reactive thymocytes during negative selection through presentation of self-epitopes by medullary thymic epithelial cells (mTECs). It is however conceivable, that weakly self-reactive thymocytes evade being weeded out if their corresponding self-antigens are presented in the thymus at levels that are too low to confer apoptosis. Once released as mature T-cell into the periphery, such clones may pose a threat, especially if they become effectively primed by means of an inherent, much stronger reactivity towards foreign pMHCs. Upon differentiation into CTLs, CD8+ T-cells may then carry out tissue-damaging effector functions if they encounter weakly cross-reactive pMHCs in considerably higher numbers, especially under inflammatory conditions associated with elevated MHCI- and MHCII-mediated antigen presentation. Increasing evidence suggests that this or a similar scenario reflects the etiology of autoimmune disorders like type 1 diabetes (T1D), a disease characterized by progressive T-cell-mediated destruction of pancreatic β-islet cells through recognition of highly abundant self-epitopes.

The mechanisms underlying antigen sensitivity, plasticity, and discrimination remain only poorly understood ^10^. This is in part because TCRs engage their corresponding pMHCs in dynamic environments, initially on antigen-scanning microvilli ^11^ and later within the immunological synapse, where interaction partners become subject to molecular crowding and mechanical forces (reviewed in ^12,13^). Recognition dynamics are hence influenced not only by intrinsic biochemical properties of the interaction partners but also by cell biological and physiological factors, which are not accounted for by conventional biochemistry and classical, equilibrium-based thermodynamics ^14^.

Furthermore, sensitized T-cell antigen detection is in most cases strictly dependent on CD8- or CD4-coreceptor binding to pMHC, and arguably to the same pMHC ligand the TCR is ligated to. Supported by structural considerations ^15^, such a scenario would position the TCR/CD3 complex and the coreceptor-associated Lck in close enough proximity to the TCR/CD3 complex for ITAM-phosphorylation. Yet, MHC-interactions are barely detectable for CD4 ^16^ and of low affinity for CD8 ranging from 100 to 1550 µM (on average 145 µM) when measured *in vitro* ^17–20^. This raises the question as to what extent tri-molecular synaptic pMHC-TCR-coreceptor complexes can be formed. Alternatively, and based on the interpretation of mechanical assays involving the use of biomembrane force-probes, CD8-MHC engagement has been proposed to promote the formation of so-termed dynamic catch bonds between cognate TCRs and their nominal pMHCs but not bystander pMHCs ^21^. Unlike slip-bonds, which account for most protein-protein interactions as they suffer in stability from applied forces, catch-bonds gain by definition in strength when placed under mechanical strain ^22^. Regardless of the exact mechanisms involved, the degree to which CD8-co-engagement enhances autorecognition has in fact not yet been determined in primary CTLs, and hence the role of CD8 in antigen discrimination remains unclear.

In this study we set out to define molecular determinants governing highly sensitized antigen detection against a background of abundant endogenous and structurally similar antigens. To this end, we mapped the permissive range of CD8+ T-cell antigen recognition dynamics in settings of anti-viral immunity and cross-reactive autoimmunity. For quantitative and molecular analyses, we devised non-invasive live cell fluorescence imaging with single-molecule resolution, preserving in such fashion cellular context essential for meaningful analysis.

Of note, virus-specific T-cells expressing the public RA14 TCR specific for a CMV epitope (phosphoprotein 65 -derived peptide presented in the context of HLA-A*0201, A2, A2/CMV) ^23,24^ responded to a single antigen, however, in a fashion that was strictly CD8-dependent with a 1000-fold reduction in sensitivity in the absence of CD8:MHCI binding. When triggered by high-quality antigens present in low abundance, a few fully phosphorylated TCR/CD3 complexes with ensuing proficient ZAP70-activation appeared sufficient to robustly stimulate T-cells. In contrast, T1D-associated T-cells which had been equipped with the 1E6 TCR and which are cross-reactive to the HLA-A*0201 presenting preproinsulin peptide (A2/PPI) ^25,26^, required on average ∼ 4000 times as much antigen for activation in a fashion that was only moderately affected by CD8-co-engagement. This enormous sensitivity gap was associated with more than 10-times shorter lifetimes of synaptic TCR:pMHC complexes (200 milliseconds vs. three seconds), as determined with the use of a newly devised single molecule FRET-based assay. In turn, such short-lived TCR:pMHC interactions supported only inefficient ZAP70 activation due to incomplete CD3-ITAM-phosphorylation.

Taken together, these observations strongly support the notion that the degree of ITAM-phosphorylation at the level of a few individual ligand-engaged TCR/CD3 complexes rather than the total number of ITAM-phosphorylation events controls the degree of ZAP70 activation as a rheostat for TCR-mediated signaling. CD8-co-engagement, which stabilized weak and more stable TCR:pMHC interactions to a similar if not equal extent, enhanced selectively virus-specific recognition by RA14 T-cells, an observation which implicates an essential role co-receptors in antigen discrimination by T-cells.

In conclusion, our findings advance our comprehension of T-cell antigen recognition and discrimination within the framework of CD8-conditional efficacy of TCR-signaling, TCR-cross-reactivity, and the high abundance of endogenous pMHCs.

## RESULTS

### Gauging human T-cell antigen sensitivity with the use of orthotopically TCR-exchanged primary human T-cells interacting with protein-functionalized planar glass-supported lipid bilayer (SLB)-based platform

We wished to compare viral and self-epitopes with regard to their recognition and cross-recognition, respectively, by CD8+ T-cells, since they represent two extremes on opposite ends within the spectrum of antigens that can in principle still elicit productive TCR-signaling. To this end, we first set out to quantitate the number of antigens required for effective T-cell activation by single-molecule fluorescence microscopy. Such endeavors are impeded by the three-dimensional nature of T-cell-target cell interactions, high cellular background, as well as phototoxicity and excessive photobleaching. Furthermore, direct imaging of specific pMHCs on the surface of living APCs is beyond realization since loading surface-exposed MHC molecules with fluorescent peptides gives rise to forbidding non-specific fluorescence background due to cellular MHC-independent internalization of the peptide. Furthermore, specific fluorophore-labeling of antigenic peptides invariably affects either presentation via MHCI or recognition by T-cells. This is because the binding cleft of MHCI is closed towards both ends, which prohibits the use of a short C- or N-terminal tethers for peptide-dye-linkage. Moreover, direct fluorescence-tagging of the original 8-10mer peptide is in most cases unproductive, as this would directly affect peptide-MHC or peptide-TCR binding.

To sidestep these issues altogether, we devised an imaging procedure in which glass-supported lipid bilayers (SLB) are functionalized with recombinant proteins of choice to serve as a surrogate target cell for the recognition by scanning cytolytic T-lymphocytes (CTLs) (**Figure 1A**). More specifically, we produced extracellular portions of peptide-loaded A2, ICAM-1 for cell adhesion via the integrin LFA-1 and B7-1 for co-stimulation by means of CD28 (**Figure 1B**). For SLB-anchorage, these proteins featured a C-terminal poly-histidine (His_12_) tag which interacted with the synthetic lipid Ni-NTA DGS present in the SLB. As verified by Fluorescence Recovery after Photobleaching (FRAP), 80% to 95% of all fluorochrome-conjugated proteins were laterally mobile when embedded in SLBs (**Figure 1C**).

**Figure 1.**
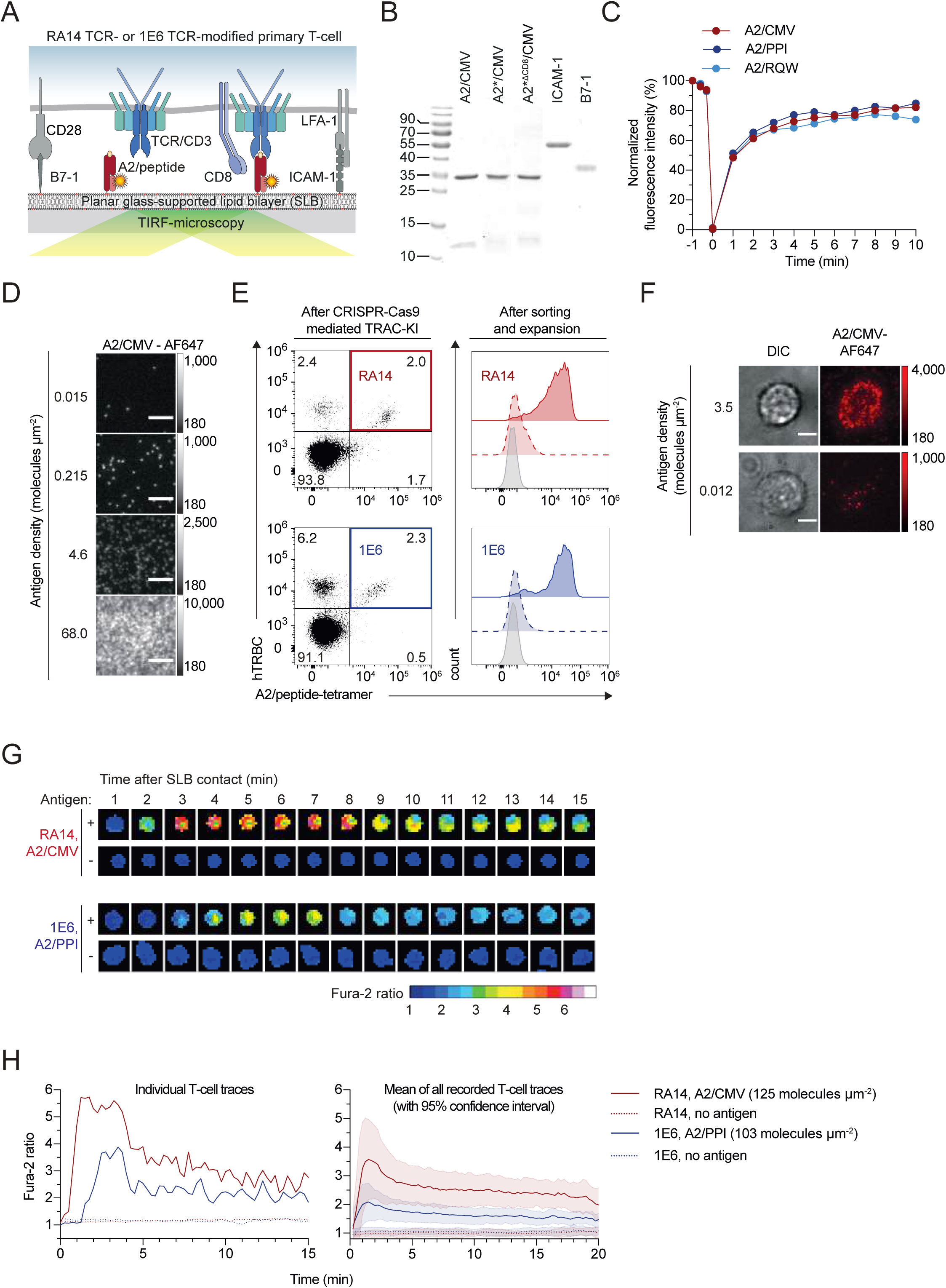
Protein-functionalized glass-supported lipid bilayers (SLB) to serve as platform for quantitative and highly resolved imaging. (**A**) Schematic representation of the SLB system used in this study. For T-cell stimulation and synapse formation, SLBs were equipped with fluorescently labeled A2 (A2/peptide) as well as ICAM-1 adhesion molecules and B7-1 co-stimulatory proteins. Extracellular domains were expressed as recombinant proteins featuring a 12x histidine tag (12xHis) for SLB-attachment via NiNTA-DGS present in the SLB at 2%. (**B**) 12.5% SDS PAGE of recombinant proteins (i.e. their c-terminally polyhistidine-tagged extracellular domains) for SLB functionalization run under reducing and boiling conditions. A2 refers to unlabeled HLA-A*0201, A2* to fluorescence-labeled HLA-A*0201 and A2*^ΔCD8^ to HLA-A*0201 mutated at position DT227/8KA to abrogate binding to CD8. (**C**) Lateral mobility of proteins on SLBs as shown by Fluorescence Recovery after Photobleaching (FRAP). Fluorescence intensity was normalized to initial intensities and plotted over time. (**D**) Representative micrographs of SLBs featuring AF647-labeled A2 at indicated densities (0.015 to 68 molecules µm^-2^). Scale bar, 5µm. (**E**) Staining of RA14 (red) and 1E6 (blue) TCRs in primary human CD8+ T-cells 5 days after CRISPR-Cas9-mediated orthotopic TCR-knockout and TCRα constant-knock-in (TRAC-KI) in primary human CD8+ T-cells. RA14 and 1E6 T-cells were isolated by staining of the human TCR β-chain (hTCRB) with TCR α/β - reactive antibody (clone: IP26, Alexa Fluor 647) and A2/peptide-specific tetramer staining (**left panel**). Specificity of A2/peptide tetramer-staining of FACS-sorted RA14 (red) and 1E6 (blue) TCR T-cells, which had been expanded in an antigen-specific fashion, with saturating amounts of nominal (full line) and irrelevant (dotted line) pMHC-tetramers. Unstained cells are shown in grey (**right panel**). (**F**) Ligand recruitment by RA14 and 1E6 T-cells contacting SLBs featuring nominal antigen at indicated densities. DIC = Differential Interference Contrast; A2/CMV-647 = Alexa Fluor 647 fluorescence image. Diffraction limited spots in the lower right panel (antigen density = 0.012 molecules µm^-2^) indicate individual fluorophores. (**G**) Time-lapse of a single Fura-2-labeled RA14 and 1E6 T-cells shows changes in intracellular calcium levels upon contact formation of SLB featuring ICAM-1, B7-1 and antigen. Temporal changes in normalized Fura-2 ratios (G) are indicated for the depicted RA14 or 1E6 T-cells. (**H**) Fura-2 ratios measured over time for the RA14 and 1E6 T-cells shown in (G, **left panel**). Median Fura-2 levels of T-cells contacting SLBs functionalized with ICAM-1, B7-1 as well as A2/CMV and A2/PPI_15-24_, respectively, at indicated densities (**right panel**). T-cells fail to flux calcium when confronted with SLBs featuring ICAM-1 and B7-1 but no antigen (n ≥ 200 cells for each condition depicted, representative of n=3). 95% confidence intervals are shaded.

The use of SLBs afforded full control over the number of antigens and accessory molecules presented to the T-cells (**Figure 1D**), which, when equipped with a matching TCR (**Figure 1E**), readily formed immunological synapses when encountering antigen on the SLB (**Figure 1F**). It also enabled the application of total internal reflection (TIRF) imaging modalities with concomitant reduction in background fluorescence due to the low depth of light penetration (<100 nm) into the imaged T-cell sample. Together, this allowed us to precisely quantitate the number of SLB-resident T-cell ligands outside (**Figure 1D**) and within forming T-cell synapses (**Figure 1F**) and to correlate their numbers with the signaling response of SLB-engaged T-cells involving the mobilization intracellular calcium as a second messenger (see below and **Figure 1G, H**).

For defined measurements and comprehensive data analysis we opted to employ primary human CD8+ T-cells featuring a defined TCR rather than antigen-specific yet polyclonal TCRs. To this end we orthotopically replaced endogenous TCRs in primary human central memory CD8+ T-cells via a CRISPR-Cas9-based methodology (for details please refer to the Methods section) with a TCR of choice ^27^. For our studies we selected (i) the virus-specific A2/CMV (pp65)-reactive public RA14 TCR and (ii) the T1D-associated autoreactive 1E6 TCR, which is specific for HLA-A*0201 (A2) complexed with the preproinsulin peptide (residues 15-24: ALWGPDPAAA, A2/PPI). As shown in Figure 1E, we achieved successful insertion of the TCR genes of interest into the TCRα locus and sorted TCR-exchanged T-cells based on co-staining of TCR-reactive IP26 antibody and PPI- or CMV-specific tetramers (left panel). We confirmed subsequent antigen-specific expansion of exclusively TCR-edited T-cells by specific tetramer staining (right panel).

To gauge the functional response of CMV-specific RA14 and self-reactive 1E6 T-cells, we employed the ratiometric Fura-2-dye to measure the intracellular calcium mobilization as a second messenger which is essential for and precedes all T-cell effector functions following antigenic stimulation. Both RA14 and 1E6 T-cells exhibited augmented calcium levels in an antigen-specific manner when confronted with SLB featuring their nominal antigens. Within minutes of initial contact with SLBs, T-cells maintained an elevated plateau of calcium influx following the peak, although this phenomenon was less pronounced in 1E6 T-cells encountering A2/PPI. Overall, the signaling response exhibited by both virus-specific and autoreactive T-cells underscores the robustness of the SLB-based imaging platform. Furthermore, the use of SLBs as an APC surrogate revealed initial disparities in the stimulatory efficacy of viral and self-peptides on their respective T-cells (**Figure 1G, H**).

### Autorecognition by 1E6 T-cells is 4000-times less efficient than CMV-recognition by RA14 T-cells

CD8+T-cells can in principle detect the presence of a single agonist pMHC ligand, a capacity which appears to strictly depend on the quality of the cognate TCR:pMHC interaction. Typically, virus-specific CD8+T-cells exhibit a high level of sensitivity towards their respective antigens ^28,29,30^, which are distinct in primary structure when compared to highly abundant endogenous self-peptides, and tend to elicit a vigorous polyclonal CD8+T-cell-response. In contrast and due to negative thymic selection, T-cell responses against self-antigens are rare. However, the sensitivity of autoreactive T-cells which have bypassed negative selection towards autoantigen for yet unknown reasons has so far not been determined. We expected it to be inferior in view of central tolerance mechanisms, which only allow for the thymic egress of T-cells with substantially reduced or no TCR-affinities towards self-antigens.

To compare antigen detection capacities between antiviral RA14 and autoreactive 1E6 T-cells in a quantitative fashion, we confronted T-cells with SLBs featuring ICAM-1, B7-1 and titrated densities of A2/CMV and A2/PPI, respectively, and recorded the dynamics of the ensuing calcium response at 37°C at the single cell level with the use of the ratiometric calcium-sensitive Fura-2 dye. We tracked Fura-2 loaded T-cells for each condition using a particle tracking algorithm ^31^. Cells confronted with antigen-free SLBs served as negative control and for Fura-2 ratio normalization (please refer to Methods section). Acquired data were then compiled as histograms (**Figure 2A**) to indicate the measured distribution of normalized single cell Fura-2 ratio values as a function of employed antigen densities. Based on the demarcation line separating non-activated from activated T-cells, we replotted histograms as dose-response curves to indicate the fraction of activated T-cells as a function of antigen density (**Figure 2B**, **left panel**). Furthermore, to assess the quality of the calcium flux we plotted the mean single cell Fura-2 ratio recorded for the activated T-cells within the respective seeded T-cell population (**Figure 2B**, **right panel**, for more details refer to the Methods section).

**Figure 2.**
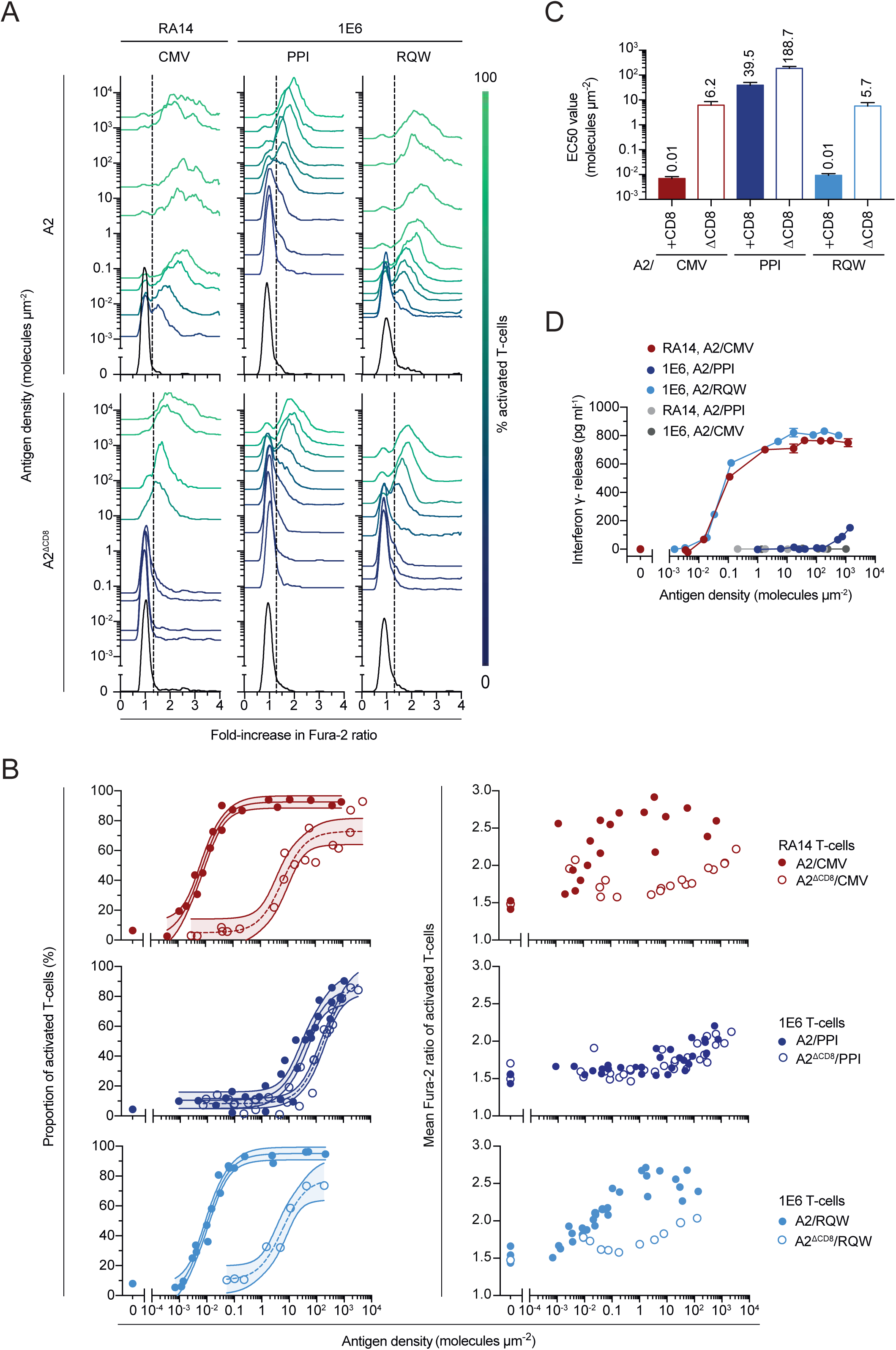
RA14 and 1E6 T-cell responses expose the wide dynamic range in T-cell antigen recognition. (**A**) Histograms illustrating the calcium response of Fura-2-labeled RA14 or 1E6 T-cells contacting ICAM-1- and B7-1-decorated SLBs presenting A2/peptide antigens in a titrated fashion. The fold-increase in Fura-2 ratio (indicative of activation) of the T-cell population is plotted against the density of antigen on the SLB. TCR-clones and peptides bound to A2 or a version of A2 (A2^ΔCD8^), which is mutated at position 227 (DT227/8KA) and no longer allows for CD8 binding, are indicated. The color of the histograms indicates the percentage of activated T-cells. Data are shown as representative of n=1-3 experiments. (**B**) Calcium response of Fura-2-labeled RA14 or 1E6 T-cells plotted as percentage of fluxing cells to A2/CMV, A2/PPI, A2/RQW as well as their corresponding CD8-binding-deficient mutants, as indicated. Negative controls represent calcium responses of T-cells facing antigen-free SLBs decorated with ICAM-1 and B7-1 molecules only (**left panel**). Mean Fura-2 ratio values of activating populations of RA14 or 1E6 T-cells are plotted against antigen densities on SLBs (**right panel**). Filled (empty) circles represent stimulation with A2 (A2^ΔCD8^). Data of n=1-3 experiments plotted. (**C**) Antigen thresholds as expressed as EC50 values after best-fit analysis of RA14 and 1E6 T-cells. (Data shown of n=1-3 replicates, error bar = s.e.m.) (**D**) Interferon-γ release by T-cells featuring indicated TCRs and facing indicated SLB-embedded antigens at indicated densities. Data points in dark and light grey represent T-cell responses towards irrelevant peptides. (Data shown of n=1-3 replicates, error bar = s.e.m.)

We found RA14 T-cells to be exquisitely sensitive towards A2/CMV, as half of them responded already to densities of 0.01 antigens per square micron (activated cells), which corresponded to 1-2 antigens per synapse. In contrast, autoreactive 1E6 T-cells required 4000-times as many copies of antigen: 50% of the SLB-engaged T-cells became activated at densities of about 40 A2/PPI molecules per square micron. Importantly, inefficient antigen detection by 1E6 T-cells did not result from T-cell- or TCR-intrinsic factors, as substituting PPI for the agonistic altered peptide ligand RQW ^32^ gave rise to reactivities similar or equal to those of RA14 T-cells confronted with A2/CMV. Of note, the much-reduced sensitivity of 1E6 T-cells towards A2/PPI was also associated with a considerably lower amplitude of the calcium response, even at antigen densities well above the activation threshold (**Figure 2B**, **right panel**). This amplitude shifted to levels we had observed for A2/CMV-engaged RA14 T-cells when we confronted 1E6 T-cells with the A2/RQW agonist (**Figure 2 A-C**).

We hence conclude that autorecognition by 1E6 T-cells was distinct from viral recognition by RA14 T-cells by quantitative and qualitative means in a fashion that depended solely on the quality of TCR:pMHC interaction. Quantitation of interferon-γ (IFN-γ) secreted by T-cells in contact with SLBs (**Figure 2D**), reflected these observed differences in the calcium response very well and testified to the validity of the SLB-based readout, which was also supported by the T-cells’ divergent cytolytic responses towards peptide-pulsed target cells (**Figure S1**). The latter, did, however, not allow for quantitative comparisons between different antigen titrations given uncertainties inherent to A2-peptide loading on the cell surface.

### Sensitized recognition of A2/CMV and A2/RQW but not autorecognition of A2/PPI is strictly dependent on CD8-engagement

Simultaneous CD8 coreceptor engagement of a non-polymorphic region of MHCI is known to sensitize T-cells for antigen, yet to a varying degree ^28,33^. Mechanisms underlying CD8-mediated sensitization are not fully understood, and current models involve (i) the recruitment of the TCR-proximal kinase Lck, which is tethered to the cytoplasmic tail of CD8, to the ligand-engaged TCR via CD8 binding of the same pMHC ^15,19^, (ii) the stabilization of the TCR:pMHC interaction via CD8-coengagement ^34–36^ and (iii) the transition from slip-bonding of the TCR:pMHC bipartite to a catch-bonding state of the tripartite TCR:pMHC-CD8 structure under TCR-imposed forces ^21,37^.

To measure the contribution of CD8 to the recognition of A2/CMV, A2/PPI and A2/RQW by RA14 and 1E6 T-cells, respectively, we abrogated CD8’s binding to SLB-embedded A2 via two mutations within the CD8-docking site (DT227/8KA, termed A2^ΔCD8^). In fact, and as already shown in **Figure 2A**, **B**, we observed a ∼1000-fold reduction in antigen sensitivity for RA14 T-cells facing A2^ΔCD8^/CMV, and in line with this, still a ∼600-fold decrease in sensitivity for 1E6 T-cells confronted with A2^ΔCD8^/RQW. In contrast, recognition of A2/PPI self-epitope by 1E6 T-cells was only 4.7-fold affected by a loss of CD8-binding.

Of note, depriving T-cells of CD8-MHCI interactions significantly reduced the amplitude of calcium response following both agonist interactions (RA14 TCR: A2/CMV; 1E6 TCR: A2/ A2/RQW) but not the less stimulatory autoimmune 1E6 TCR: A2/PPI interaction (**Figure 2B**, **right panel**). Our findings suggest that TCR:pMHC interactions have to meet kinetic as well as biophysical criteria for CD8-mediated boosting to take effect. These may include a permissive TCR:pMHC dwell time for CD8 to interact with the same pMHC ^38^ and to carry along CD8-tethered Lck for the phosphorylation of nearby ITAMs or pITAM-recruited ZAP0-phosphorylation to occur ^26^. In both agonist recognition scenarios, the reduced amplitude of calcium signaling in the absence of CD8-binding could not be compensated for by saturating antigen densities. This finding - together with the observation that CD8 failed to boost autorecognition - indicated that CD8-pMHCI-binding constitutes an integral part of the mechanisms underlying sensitized antigen recognition and efficient signal transduction but not weak cross-reactivity (see below). We hence considered it plausible that CD8-co-engagement supports the discrimination of high and lower quality T-cell antigens.

### Inefficient autorecognition by 1E6 T-cells coincides with weak antigen engagement

To explore mechanisms underlying the observed differences in antigen recognition between RA14 and 1E6 T-cells, we focused on the strength of TCR-antigen engagement within the immunological synapse as a comparative parameter. Measurements via SPR had already revealed *in vitro* substantial differences in TCR:pMHC affinities ^17,24^. Considering that synaptic ligand engagement is an active process far from equilibrium which involves the application of mechanical force, we abstained from pursuing TCR:pMHC binding constants *in situ*. Instead, as a meaningful measure for synaptic antigen engagement we determined the number of otherwise laterally mobile SLB-tethered TCR-bound pMHCs within the immunological synapse for a total of eight binding scenarios in the presence and absence of CD8-binding (**Figure 3A-C**): The RA14 TCR binding to A2(ΔCD8)/CMV as well as the 1E6 TCR binding to the autoantigen A2(ΔCD8)/PPI, the altered peptide ligand A2(ΔCD8)/RQW and to mismatched A2(ΔCD8)/CMV, which served as a negative control to assess (i) nonspecific as well as (ii) CD8-dependent retention. As a direct measure for synaptic binding strength, we determined the level of TCR-specific ligand-enrichment for various pMHC densities (**Figure 3B**). This approach was chosen due to the considerably lower lateral diffusion of TCR-bound pMHCs compared to that of free SLB-resident pMHCs, as previously observed ^39,40^, and the ensuing enrichment of retained TCR-bound pMHCs. Sizes of immunological synapses forming with antigen-functionalized SLBs were comparable (**Figure 3C**), yet contacts of T-cells with SLBs featuring irrelevant pMHCs (1E6 T-cells confronted with A2/CMV) were significantly less stationary.

**Figure 3.**
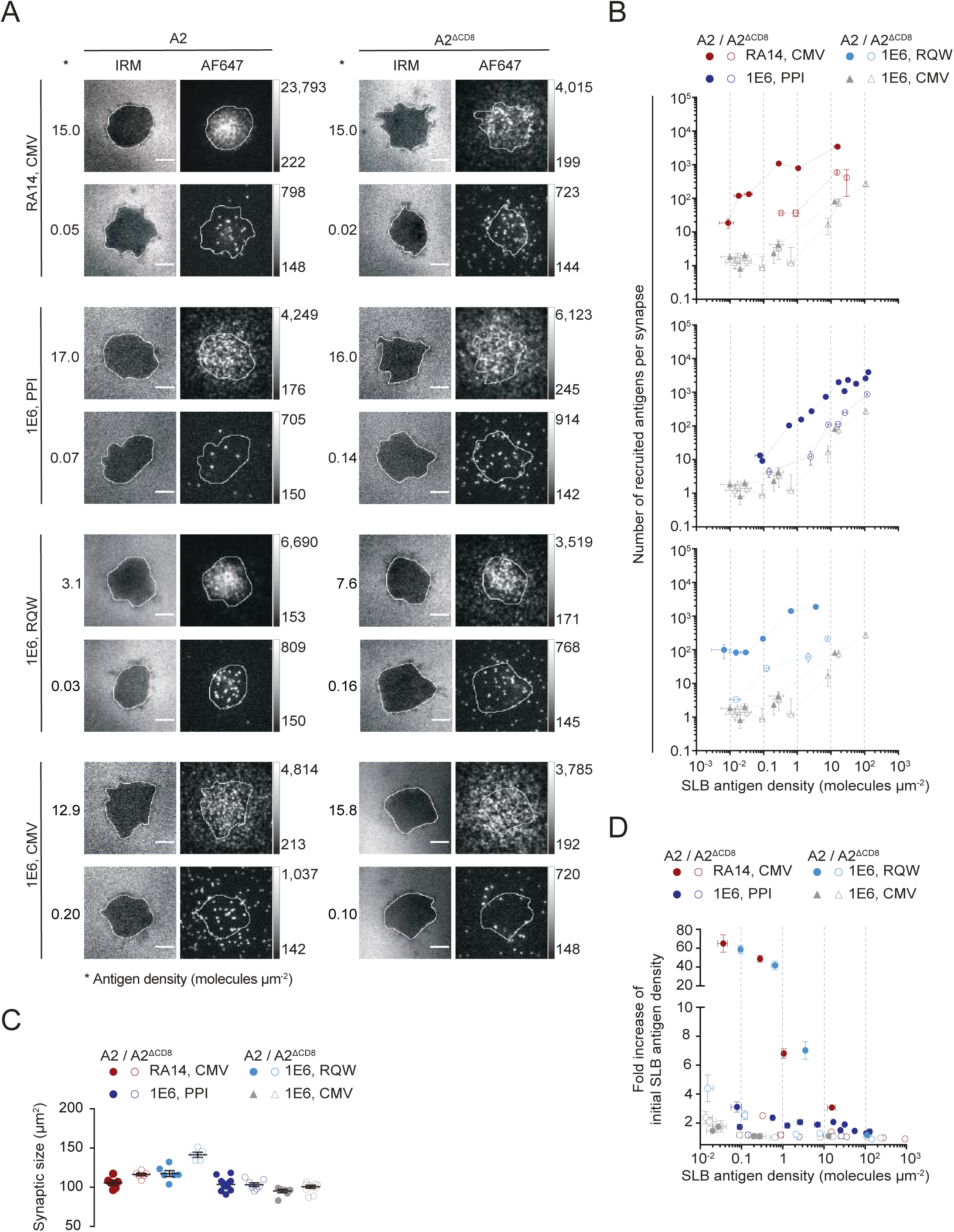
pMHC-recruitment-based assay reveals considerable differences in synaptic TCR:pMHC binding strength and the influence of CD8 engagement. (**A**) Micrographs of primary T-cells featuring the RA14 or 1E6 TCR and facing SLB-resident antigens (A2/CMV, A2/PPI, A2, RQW) at indicated densities. Interference Reflection Microscopy (IRM) indicates areas of close T-cell-contacts with the SLB. Note the degree of accumulation of AF647-labeled A2/peptide complexes in the AF647 channel. Scale bar, 5 µm. (**B**) Numbers of pMHC ligands accumulating in synapses of indicated T-cells (Y-axis) are plotted against the density of pMHCs present on the SLB (X-axis). The type of TCRs, pMHC ligands and versions of MHCI (A2, A2^ΔCD8^) are indicated. (n=1, with 15-35 cells per condition, error bars = s.e.m.) (**C**) Size of synaptic area as deduced via IRM is plotted against indicated T-cells confronted with indicated antigens. A2/CMV and A2^ΔCD8^/CMV (grey triangles) serve as negative controls for 1E6 T-cells to determine the degree of non-specific as well as CD8-dependent (yet TCR-independent) ligand accumulation. Error bars = s.e.m. (**D**) Fold-increases of initial SLB antigen densities are plotted against initial antigen densities on the SLB. (n=1, with 15-35 cells per condition, error bars = s.e.m).

For reliable assessments we first recorded the level of non-specific and purely CD8-mediated enrichment. As shown in **Figure 3 A, B** such enrichments were minimal and indistinguishable between the use of both mismatched A2/CMV and A2^ΔCD8^/CMV (1.05), (i) testifying to the integrity of the SLB-based scoring system and (ii) also revealing only a neglectable contribution of CD8-MHC binding to overall synaptic ligand enrichment.

In contrast, synaptic binding of both the RA14 TCR and the 1E6 TCR to their agonist ligands A2/CMV and A2/RQW, respectively, was considerable and gave rise to a three- to 65-fold ligand enrichment, depending on the number of available pMHC ligands (**Figure 3D**). Of note, both binding scenarios were unlike that observed for mismatched TCR:pMHCI pairs highly dependent on CD8-engagement with a more than tenfold loss in synaptic binding after abrogation of CD8-binding. Since CD8-binding alone, i.e. synaptic pMHCI enrichment in the presence of mismatched TCRs, did not give rise to measurable values, this observation supported the notion of cooperative pMHC-binding between the TCR and CD8.

In line with the rather moderate levels of T-cell recognition, we recorded significantly decreased levels of synaptic binding between the 1E6 TCR and the self-ligand A2/PPI. Especially at lower antigen densities, i.e. when TCRs were not outnumbered by SLB-resident pMHCs, enrichment was ten-to 25-times reduced when compared to that of both agonist TCR:pMHC pairings (**Figure 3D**). Of note and supportive of cooperative pMHC-binding between the TCR and CD8, abrogating CD8-binding reduced TCR:pMHC binding to levels which could no longer be distinguished from non-specific binding. However, despite minimal binding, i.e. below the detection level of the recruitment-based assay, A2^ΔCD8^/PPI proved unlike the unmatched negative control (A2^ΔCD8^/CMV) still stimulatory at densities higher than 100 molecules per square micron (**Figure 2A-C**).

Taken together, our results reveal significant disparities in synaptic TCR:pMHC binding strength between TCR interactions with genuine agonist ligands and an autoreactive ligand, which strongly correlate with their respective stimulatory capacities. Notably, we did not detect any CD8-pMHC binding in the absence of a cognate TCR. However, we observed a substantial contribution of CD8 to the overall TCR:pMHC binding strength for all three TCR:pMHC pairings, implying cooperative pMHC-binding mechanisms between the TCR and CD8. This cooperative interaction remained evident in the case of the 1E6 TCR:A2/PPI pairing even at pMHC densities below 0.3 molecules per square micron - well below the activation threshold. This finding points to factors other than activation-induced changes within synaptic architecture which contribute to altered, CD8-dependent TCR:pMHCI engagement behavior.

### Förster Resonance Energy Transfer (FRET)-based imaging visualizes synaptic TCR:pMHC binding dynamics for quantitative analysis

To comprehensively elucidate the determinants of differential antigen sensitivity and CD8-dependency at the molecular level, we set out to delineate the kinetics of synaptic TCR:pMHC binding. While surface plasmon resonance (SPR)-based *in vitro* measurements faithfully capture the intrinsic binding properties of TCR and pMHC, they neither factor in geometric constraints of binding within the developing immunological synapse nor mechanical forces acting *in situ* on TCRs and their ligands.

To overcome these limitations, we developed a non-invasive imaging technique based on Förster Resonance Energy Transfer (FRET), a quantum-mechanical, radiation-free phenomenon involving dipole interactions between matching fluorescent dyes. FRET efficiency is inversely proportional to the sixth power of the inter-dye distance, turning it into a precise molecular ruler for spatial proximities in the 2 to 10 nanometer range ^39^. Here we employed FRET as a digital indicator for the presence or absence of TCR:pMHC binding.

In our approach, illustrated in **Figure 4A**, we converted the TCR/CD3 complex into a FRET donor by employing a newly engineered recombinant single-chain antibody fragment (scF_V_) derived from the CD3ε-reactive monoclonal antibody UCHT-1. We labeled this UCHT-1-scF_V_ site-specifically with Alexa Fluor 555 (AF555) by targeting a newly introduced unpaired cysteine residue with AF555-maleimide. To serve as a FRET acceptor, A2/peptide complexes residing on the supported lipid bilayer (SLB) were site-specifically labeled with Alexa Fluor 647 (AF647)-maleimide via a cysteine residue introduced via mutagenesis into the β2-microglobulin (β2m) subunit. According to existing cryo-electron microscopy ^41^ and X-ray ^42^ structural data, we expected this arrangement to position the two corresponding fluorophores within a proximity of approximately 2 nanometers upon TCR:pMHC binding—i.e. well below the Förster radius of around 5.1 nanometers, at which half-maximal energy transfer occurs. To arrive at optimal FRET yields and assay compatibility, we conducted extensive tests on various FRET-labeling positions within UCHT-1-scF_V_ and A2, and selected positions for cysteine substitutions within the UCHT-1-scF_V_ at position 15 (S15C) and within β2m of the MHC class I molecule at position 2 (β2m I2C), as is depicted (**Figure 4A**, **right panel**).

**Figure 4.**
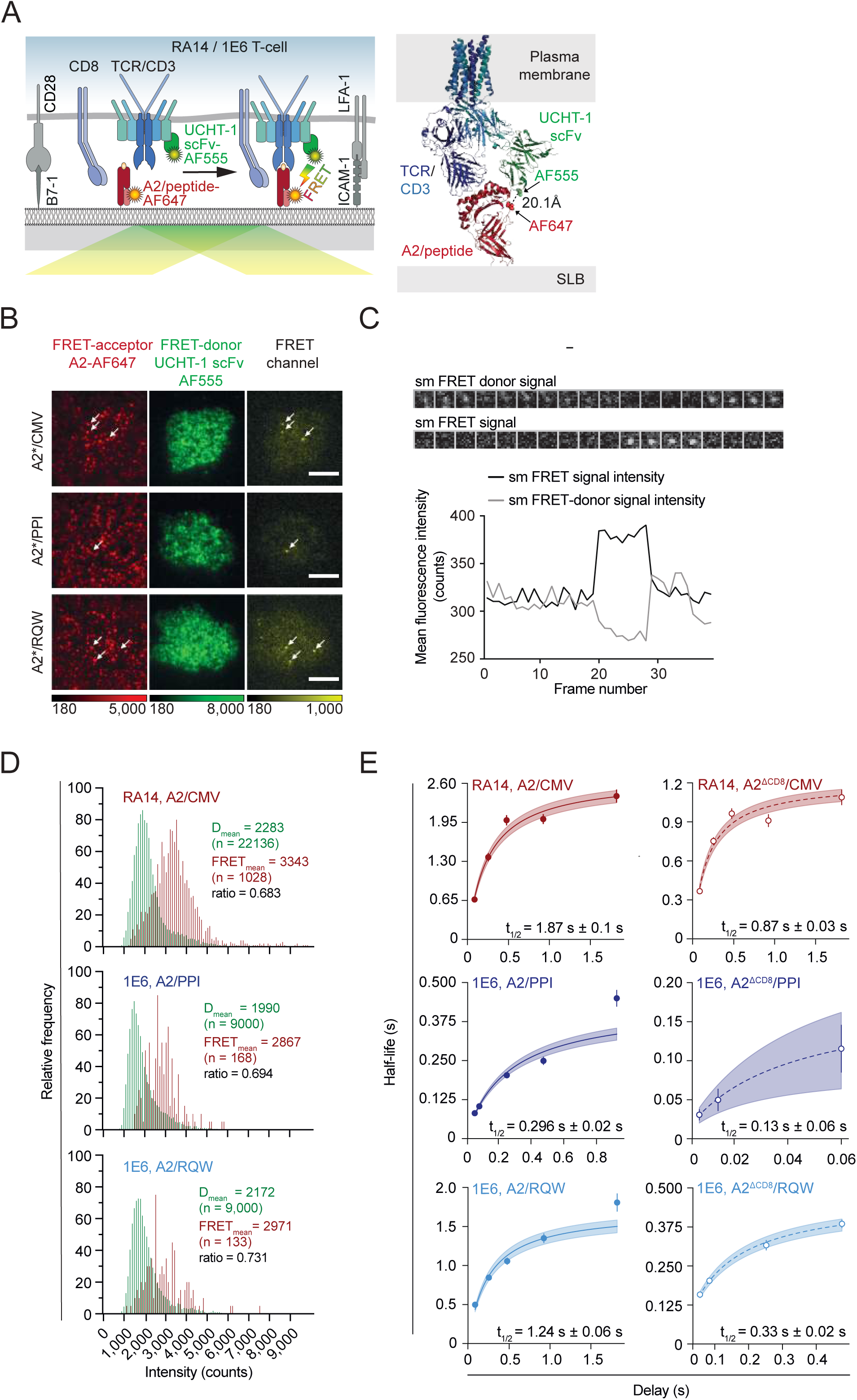
Single-molecule FRET microscopy demonstrates stabilizing impact of CD8-coreceptor engagement on TCR:pMHC interactions and reveals short lifetimes for autoreactive TCR:pMHC interactions. (**A**) Scheme depicting the FRET-based system devised to quantitate synaptic TCR-pMHC binding (**left panel**). T-cells are decorated with an anti-CD3 scF_V_ (UCHT-1 scF_V_) which has been site-specifically conjugated with Alexa Fluor 6555-maleimide via an unpaired cysteine residue in place of serine at position 15 to serve as FRET-donor. The β2m subunit of A2 is modified at position 2 with an unpaired cysteine (β2m I2C) and conjugated with AF647-maleimide to act as FRET-acceptor. The adhesion and co-stimulation are provided by ICAM-1 and B7-1, respectively. Structural model of the TCR/CD3 complex (blue) decorated with UCHT-1 scF_V_ (green) binding to peptide/A2 (red, **right panel**). Positions conjugated with FRET acceptor and FRET donor dyes are indicated as green spheres on UCHT-1 scF_V_ and red spheres on A2, respectively, and give rise to a predicted inter-dye distance of ∼20 Å. |(**B**) FRET acceptor, FRET donor and FRET images displaying synapses of RA14 and 1E6 T-cells in contact with SLBs which presented ICAM-1, B7-1 as well as indicated antigens. Discernible diffraction-limited single-molecule FRET events (white arrows) define binding events between individual synaptic TCRs and A2/peptide ligands. (**C**) Representative trace (**top**) and corresponding fluorescence intensities (**bottom**) of single-molecule FRET event and its corresponding FRET donor event recorded over time within the synapse of an RA14 T-cell in contact with an SLB featuring A2/CMV (time interval: 252 milliseconds). Event appeared and vanished within one frame. (**D**) Histograms summarizing intensities of single UCHT-1 scF_V_-AF555 (green) and single-molecule FRET events (red) recorded in TIRF mode as indicated in synapses of SLB-engaged RA14 and 1E6 T-cells scanning for A2/CMV, A2/PPI or A2/RQW. The ratio built from average single-molecule intensities was derived from diffraction-limited UCHT-1 scF_V_-AF555 and FRET events, is specific for the FRET- and microscopy system employed. Data of n=2 experiments shown. (**E**) Non-linear least squares regression fit of measured lifetimes (37°C) of synaptic TCR-peptide/HLA interactions recorded for RA14 and 1E6 T-cells scanning SLBs for indicated pMHCs. Time delays between recorded images are as indicated (7 ms, 10 ms, 14 ms, 28 ms, 56 ms, 84 ms, 252 ms, 476 ms, 924 ms, 1820 ms). Shaded areas refer to 95% confidence intervals. Measured half-lives and SDev are indicated. Data shown as representative of n=2 independent experiments. For more details, please refer to Methods section.

Decoration of T-cells with UCHT-1 scF_V_ was specific and yielded even higher staining intensities when compared to that achieved with the use of the TCR-reactive IP26 monoclonal antibody (**Figure S2A**, **B**). We consider it likely that given the smaller size of scF_V_s a greater number of surface -exposed CD3 epitopes become accessible. Quantitative analysis of UCHT-1-scF_V_ and IP26 antibody surface densities indicated similar numbers of surface-accessible TCR/CD3 complexes, approximately 40 receptors per square micron (**Figure S2C, D**). This further confirmed similar, if not identical, TCR surface expression levels between TCR-knocked-in T-cells and non-modified CMV-specific T-cells isolated through pMHC-enrichment from the blood of CMV- and A2-positive donors prior to peptide-dependent expansion.

We subsequently assessed whether the integration of fluorophores into β2m (I2C) or the coating of T-cells with the UCHT-1-scF_V_ probe affected T-cell antigen recognition in our SLB-based system. In contrast to the results obtained with the use of full UCHT-1 antibody exposure ^43^, the decoration of T-cells with UCHT-1-scF_V_ activate T-cells in the absence of antigen (**Figure S2E**, upper panel). Moreover, it did not exhibit any discernible impact on T-cell sensitivity to antigen, as assessed by monitoring intracellular calcium mobilization in RA14 and 1E6 T-cells interacting with protein-functionalized SLBs (**Figure S2E**, lower panel).

Only when UCHT-1-scF_V_-decorated RA14 and 1E6 T-cells encountered stimulatory pMHC, specific FRET-sensitized emissions were detected in the FRET channel (**Figure S3A**). As anticipated, the FRET signal promptly disappeared upon the quantitative removal of the FRET acceptor, accompanied by a significant increase in FRET donor intensity (**Figure S3A**). Correlating FRET values as measured by FRET <u>D</u>onor <u>R</u>ecovery <u>A</u>fter FRET <u>A</u>cceptor <u>P</u>hotoablation (DRAAP) with measurements of fluorescence-<u>S</u>ensitized <u>E</u>mission FRET (SE-FRET) gave rise to highly similar slopes for the three different recognition scenarios tested (1. RA14 → A2/CMV; 2. 1E6 →A2/PPI; 3. 2. 1E6 → A2/RQW), further confirming the integrity of the FRET-based approach (**Figure S3B**). In addition, FRET signals exhibited a dose-dependent increase with rising antigen densities, confirming the validity of the FRET signal as a measure for specific TCR:pMHC binding (**Figure S3C**). Consistent with above findings from synaptic recruitment experiments, RA14 T-cells engaging A2/CMV exhibited considerably higher antigen-dependent FRET values than 1E6 T-cells interacting with A2/PPI, thus confirming the considerably weaker synaptic TCR-antigen binding in the case of autorecognition (**Figure S3C**).

### A2/PPI engaged by cross-reactive 1E6 TCRs feature elevated synaptic off-rates

We next sought to visualize single-molecule FRET (smFRET) events, as tracking them over time allows for precise determination of TCR:pMHC binding lifetimes within the immunological synapse. To minimize FRET donor bleed-through into the FRET channel, we reduced the number of probe-associated TCR/CD3 complexes by employing a mixture of unlabeled and site-specifically fluorophore-conjugated UCHT-1-scF_V_. With this approach we arrived at a TCR/CD3-labeling ratio of 1:5 to 1:10, facilitating the recording of smFRET events with satisfactory signal-to-noise ratios ranging from 10:1 to 30:1. To further enhance separation of individual binding events, we labeled only 10 to 30% of the SLB-resident pMHCs with the FRET acceptor dye AF647 in a site-directed manner and adjusted them to a density of approximately 15 molecules per square micron (**Figure 4B**). To qualify as smFRET events, individual signals in the FRET channel had to meet the following criteria: they had to appear and disappear as diffraction-limited spots within a single step (**Figure 4C**) and furthermore had to give rise to a brightness distribution resembling that of single, diffraction-limited FRET donor signals (**Figure 4D**). Notably and most likely the result of similar docking geometries, the ratios between the average intensities of single UCHT-1-scF_V_-AF555 events and single-molecule FRET events were indistinguishable for all three TCR:pMHC pairs investigated (RA14 → A2/CMV, 1E6 → A2/PPI, 1E6 → A2/RQW, **Figure 4D**).

The observation that the stimulatory potency of TCR:pMHC engagement is often inversely correlated with the corresponding off-rate has led to the formulation of the kinetic proofreading model as a way to predict downstream signaling based on binding properties ^38^. To assess the relevance of this model for autorecognition we compared the synaptic off-rates for the RA14 and 1E6 TCRs with their respective SLB-embedded antigens using the single-molecule FRET-based approach described above. More specifically, we tracked single synaptic TCR:pMHC binding events as diffraction-limited single-molecule FRET events over time which appeared and disappeared in discrete steps and in a manner similar to previous quantifications of TCR:pMHC binding dynamics in murine TCR-transgenic systems ^9,44,45^. We derived photobleaching-affected apparent half-lives from individual single-molecule FRET traces by survival analysis and a weighted non-linear least squares regression fit. To arrive at the true rate of synaptic bond dissociation, we accounted for fluorophore photobleaching in a fashion that is described in detail in the Methods section.

In agreement with McKeithan’s proofreading model, the RA14 TCR interacted within the immunological synapse with the viral A2/CMV antigen with a half-life of 1.73 ± 0.07 seconds at 37°C (mean value of 2 measurements, **Figure 4E, Figure S4A**) and 5.3 ± 0.65 seconds at 26°C (**Figure S4B**). Of note, synaptic TCR:pMHC interactions appeared to be twice as long-lived, at least when compared to SPR-based measurements carried out at 25°C ^46^. In contrast, synaptic 1E6 TCR -A2/PPI interactions featured a half-life of only 323 ± 0.023 milliseconds (322 milliseconds ± 0.027 milliseconds, 26°C), in line with their (i) vastly reduced stimulatory potency (**Figure 2B**) as well as their (ii) considerably lower synaptic TCR-ligand recruitment (**Figure 3A, B**) and (iii) ensemble FRET yields (**Figure S3C**). After substituting the self-peptide PPI with the altered peptide ligand RQW, 1E6 TCR interactions increased in half-life to 1.12 ± 0.06 milliseconds (3.21 ± 0.29 seconds at 26°C), again consistent with their highly stimulatory potency, elevated synaptic recruitment and ensemble FRET yields.

In summary, with the use of our newly established smFRET-based imaging approach we precisely determined synaptic TCR:pMHC lifetimes in settings of viral and autorecognition within the millisecond range. Reflective of their 1000- to 4000-fold reduced stimulatory strength, the autoreactive 1E6 TCR-A2/PPI interactions featured five to 16-fold reduced lifetimes with predictable consequences for rather inefficient downstream signaling.

### Synaptic TCR:pMHCI lifetimes are affected by CD8-MHCI co-engagement independent of their length

When measured *in vitro*, CD8 exhibits on average an affinity towards MHCI between 100.5 µM and 115 µM (at 25°C) ^36,47^, in fact surpassing that of the 1E6 TCR (∼ 335 µM) and other autoreactive TCRs towards their respective self-peptide/MHC ligands ^26^. This contrasts our observations that CD8-MHCI engagement failed to register within the immunological synapse in the absence of TCR-specific antigens while it substantially boosted synaptic TCR:pMHC binding in all three scenarios investigated in (**Figure 3A, D, E**). As demonstrated above, CD8-engagement sensitized in a dramatic fashion the detection of the agonist ligands A2/CMV and A2/RQW by RA14 and 1E6 T cells, respectively, while it only marginally affected cross-reactive auto-recognition of A2/PPI by 1E6 T-cells.

To approach a biophysical foundation underlying these behaviors, we examined how CD8 engagement influenced the stability of synaptic TCR:pMHC binding. Cooperative pMHC-binding by the TCR and CD8 had previously been suggested to transform agonist but not weak agonist TCR:pMHC slip-bonding into TCR:pMHC:CD8 catch bonding ^21,37^. If true, CD8:pMHCI-binding would be expected to stabilize synaptic pMHC interactions with agonist-TCRs but not with weak-agonist autoreactive TCRs. To test this we quantitated the lifetimes of synaptic TCR:pMHC interactions via smFRET in the absence of CD8-pMHC binding. We observed - consistent with the stabilizing effect of CD8 - a two- to three-fold reduction in synaptic lifetimes for the RA14 TCR and 1E6 TCR engaging A2^ΔCD8^/CMV and A2^ΔCD8^/RQW, respectively (**Figure 4E, Figure S4A, B**). This was accompanied with a tenfold decrease in overall binding strength as assessed via synaptic pMHC recruitment analysis (**Figure 3B**), implying a role for CD8 in accelerating synaptic TCR:pMHC on-binding.

Interestingly, we observed a similarly stabilizing effect of CD8 on synaptic 1E6 TCR binding in the context of cross-reactivity when providing the auto-antigen PPI (**Figure 4E, Figure S4A, B**). The half-live of 1E6 T-cells binding to A2^ΔCD8^/PPI was 2.4-times more short-lived compared to that of A2/PPI at 37°C and 1.4-fold reduced when measured at 26°C. This strongly suggested a similar role of CD8 engagement on synaptic binding life times of weak, cross-reactive ligands as well as highly stimulatory agonist ligands.

Although the autoreactive 1E6:A2/PPI interaction lifetime doubled in the presence of CD8, we consider it likely that overall TCR:pMHC dwell times are too short to enable an efficient CD8-mediated boost of antigen sensitivity (**Figure 2A, B**) and recruitment (**Figure 3B**), in agreement with the kinetic proofreading model. Of note, due to the exceedingly low levels of synaptic binding between the 1E6 TCR and A2^ΔCD8^/PPI (with synaptic antigen enrichments being close to 1, **Figure 3D**) we were only able to determine an approximate gain in synaptic binding strength attributable to CD8. Accordingly, measurements of TCR:pMHC binding lifetimes for A2^ΔCD8^/PPI suffered from the limited frequency of interactions detectable under these very low affinity conditions.

Taken together, our results are in agreement with CD8 co-engagement stabilizing synaptic TCR:pMHC binding. Lifetimes of both barely and highly stimulatory TCR:pMHC interactions appeared to equally benefit from CD8 binding yet, as demonstrated above (**Figure 2A-D**), while signaling output was differentially affected. Our measurements are consistent with CD8 and TCRs binding simultaneously to the same pMHCI ligand, with conceivable consequences for downstream signaling (**Figure 2B**) due to the recruitment of CD8-tethered Lck to ligand-engaged TCR/CD3.

### A2/PPI-autoantigens trigger 1E6 TCRs inefficiently

To quantitatively assess how differences in TCR-binding strength (**Figure 3**) and kinetics (**Figure 4E, Figure S4**) impacted TCR-proximal signaling, we correlated for both RA14 and 1E6 T-cells the numbers of pMHCs engaged within the synapse with the magnitude of the resulting calcium response serving as readout for T-cell activation. Remarkably, we found that within the first 15 minutes after T-cells making SLB-contact, activation of 50% of RA14 T-cells correlated with the binding of only one to three A2/CMV antigens. In contrast, 100-200 times more TCRs needed to be engaged at any time for the activation of 1E6 T-cells via cross-reactive A2/PPI (**Figure 5A**), with mean calcium levels being significantly inferior for the activated cell fraction when compared to those of A2/CMV-activated RA14 T-cells (**Figure 2B**). Taken together, these comparisons underscore stark qualitative differences in TCR-trigger efficiencies observed between settings of viral recognition and cross-reactive autorecognition.

**Figure 5.**
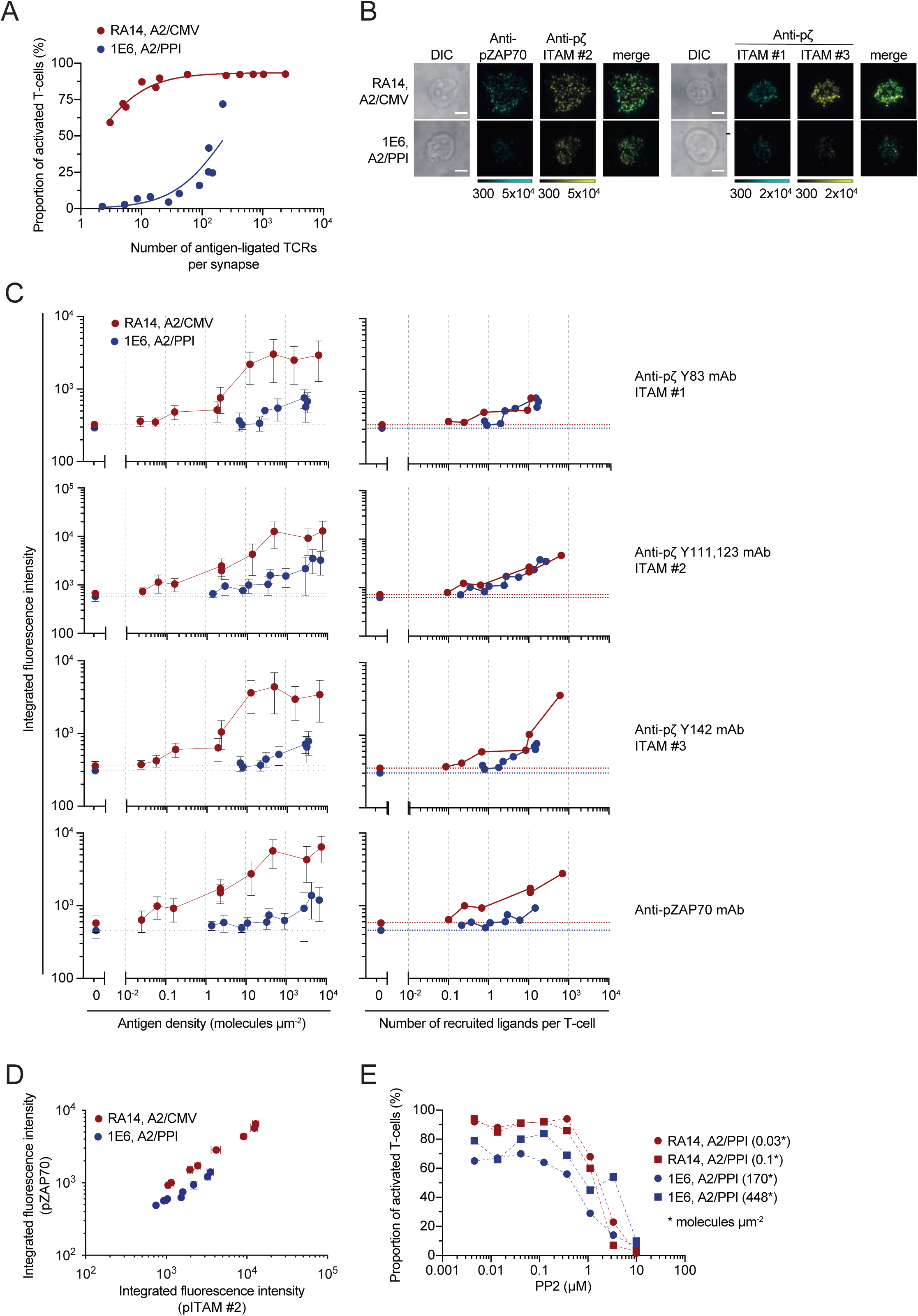
Autorecognition is characterized by inefficient ZAP70 activation in response to TCR-triggered ITAM phosphorylation. (**A**) Correlation between proportion of activated T-cells and antigen-ligated TCRs per T-cell synapse. The number of recruited ligands per synapse of RA14 and 1E6 T-cells was quantified and plotted against the proportion of activated T-cells as determined by Fura-2 ratiometric calcium imaging. The data were fit using a three-parameter dose-response curve fit (shown as solid lines). Data of n=2 experiments shown. (**B**) Quantitative immunofluorescence analysis to assess degree of ITAM-phosphorylation and ZAP70 activation. RA14 and 1E6 T-cells were seeded on SLBs featuring ICAM-1, B7-1 and, as indicated, A2/CMV and A2/PPI, and left for 12 minutes. T-cells were then fixed, membrane permeabilized and stained with antibodies targeting as indicated pITAMs and pZAP70. Immunofluorescence was recorded in TIRF mode. (**C**) Integrated fluorescence intensities associated with the use of indicated antibodies reactive against pITAMs and pZAP70 (p315) on synapses of RA14 (red) and 1E6 (blue) T-cells were plotted against indicated densities of antigen which had been placed on SLBs also featuring ICAM-1 and B7-1. T-cells confronted with ICAM-1- and B7-1-functionalized SLBs devoid of antigen served as negative controls to indicate background staining demarcated as a dotted baseline (**left panel**). Alternatively, Integrated fluorescence intensities of pITAM or pZAP70 were plotted against the number of recruited ligands per T-cell (**right panel**). Error bars = SDev; Data of one experiment shown with n = 17-46 cells per condition; scale bar, 5µm. (**D**) Synaptic T-cell-associated integrated fluorescence intensities associated with pITAM#2-reactive antibodies were plotted against integrated fluorescence intensities derived from pZAP70-reactive antibodies. Note the gap separating RA14 T-cells and 1E6 T-cells recognizing their respective antigen (A2/CMV, A2/PPI) as is indicated. Error bars = SDev; Data of one experiment shown with n = 17-46 cells per condition; scale bar, 5µm. (**E**) We plotted the calcium response of RA14 (red) and 1E6 (blue) T-cells which had been confronted with SLBs featuring ICAM-1, B7-1 and antigens in indicated densities against the concentration of the added Lck-inhibitor PP2.

We next set out to reveal the mechanisms underlying the observed deficits in cross-reactive self-recognition. For this we closely examined membrane-proximal signaling events following TCR-ligand engagement. We focused our analysis on the Lck-mediated CD3-ITAM phosphorylation as well as ensuing recruitment of ZAP70 to tyrosine-phosphorylated CD3ζ-ITAMs 1-3 (pITAM#1-#3) and its subsequent activation, two steps which determine the threshold for further downstream signaling, including the activation of phospholipase C (PLC) as a prerequisite for calcium signaling and activation-induced changes in gene transcription.

Immunostaining (**Figure 5B, C**) shows that ITAM- and ZAP70-phosphorylation increased gradually in RA14 T-cells with rising densities of SLB-resident A2/CMV. Reflecting weaker synaptic TCR:pMHC binding and the higher activation threshold, 1E6 T-cells displayed in contact with A2/PPI a similar trend though with an overall shift towards considerably higher SLB antigen densities.

Plotting ITAM-phosphorylation levels against the number of engaged pMHCs (i.e. TCR-occupancies) within synapses of individual SLB-contacting T-cells revealed that 1E6 T-cells engaging A2/PPI exhibited at all pMHC occupancies similar or only moderately lower CD3-ITAM phosphorylation levels compared to RA14 T-cells confronted with A2/CMV (**Figure 5C**).

Yet both the recruitment and activation of ZAP70, i.e. the subsequent steps following ITAM phosphorylation, were significantly reduced in A2/PPI-activated 1E6 T-cells at all pMHC occupancies (**Figure 6C**). We hence conclude that the translation of ITAM-phosphorylation into ZAP70 activation was significantly less efficient in A2/PPI-engaged 1E6 T-cells. Correlating anti-pZAP70 with anti-pITAM#2 fluorescence intensities (**Figure 5D**) further supported this notion as our results agree well with predictions based on the kinetic proofreading model of T-cell activation, which penalizes short-lived TCR-ligand interactions with a substantial loss in stimulatory potency. Quantitative analysis of our immunofluorescence data (**Figure 5C**) furthermore implies that frequent but brief TCR:pMHC interactions promoted only partial but not complete phosphorylation of a larger number of TCR/CD3 complexes, with little effect on ZAP70 recruitment and activation.

Of note, calcium signaling decreased upon titration of the Lck inhibitor PP2 to the same extent in 1E6 and RA14 T-cells facing SLBs which had been functionalized with nominal antigen in densities adjusted to activate more than 65% of the T-cells (**Figure 6E**). We therefore conclude that low-level pharmacological inhibition of Lck affected scenarios of highly sensitized and insensitive T-cell recognition equally. This finding underscores the identical nature of the underlying TCR-proximal signaling pathways, which all rely on TCR-triggered Lck kinase activity and a critical number of activated ZAP70 molecules as a starting point.

Taken together, our analysis reveals a scenario in which a few longer-lived TCR-agonist interactions induce the complete phosphorylation of a small number of signaling-competent TCRs. In contrast, short-lived TCR:pMHC interactions give rise to a higher number of partially-phosphorylated and largely signaling-incompetent TCR/CD3 complexes if they occur at high frequency and in response to high antigen levels. In agreement with this, full activation of RA14 T-cells was indeed maintained in the presence of few A2/CMV antigen (**Figure 2B**), albeit with levels of ITAM- and ZAP70-phosphorylations that fell below the detection limits of conventional immunofluorescence-based assays.

## DISCUSSION

In this study we gauged the permissive range of T-cell recognition dynamics which demarcate the boundaries of central T-cell tolerance in the specific context of (i) CD8-conditional TCR-signaling proficiency, (ii) TCR-cross-reactivity and (iii) the high abundance of structurally similar endogenous epitopes. Employing non-invasive advanced live cell imaging techniques, we focused on two distinct ends of the recognition spectrum, i.e. the detection of viral and self-epitopes ^48^. With regard to our model systems, we opted for primary human T-cells engineered with TCRs of interest via CRISPR-Cas9-mediated orthotopic TCR-exchange: (i) the public CMV-specific RA14 TCR and (ii) the autoreactive T1D-associated 1E6 TCR. As revealed in this study, the RA14 TCR conveyed ultrasensitive detection down to a single antigen in a strictly CD8-dependent manner. In contrast, the highly cross-reactive 1E6 T-cell clone required the presence of at least 4000 self-antigens for activation due to the exceptionally low *in situ* TCR-affinity, as witnessed in this study, and importantly, with little involvement of CD8. As discussed in more detail below, this behavior is best explained with CD8 playing an active part in ligand discrimination. Notably, when experimentally adjusted for equal TCR-antigen occupancies rather than antigen densities, both recognition scenarios exhibited comparable levels of CD3-ITAM-phosphorylation - representing the initial intracellular biochemical event post TCR-engagement - at the ensemble level. However, downstream signaling diverged considerably at the level of ZAP70-recruitment and activation, which explains our observation that autoreactive 1E6 T-cells required at least 100 as many self-epitope-engaged TCRs at any time for activation compared to RA14 T-cells facing A2/CMV.

For reasons outlined above we based our conclusions predominantly on the use of SLBs, which served as a quantitative and highly defined reconstructive surrogate for APCs and enabled the use of noise-reduced TIRF-microscopy as an imaging modality deemed critical for achieving single molecule precision. Testifying to the integrity of our imaging platform, virus-specific T-cells became activated in the presence of a single antigen. As demonstrated above, the use of SLBs allowed us to reveal the contribution of coreceptor-binding to antigen sensitivity as well as how it affects the stability and frequency of synaptic pMHC-TCR interactions. To assess the latter, we resorted to a newly devised single-molecule FRET-based microscopy approach, in which we employed recombinant SLB-resident peptide-loaded A2 and decorated surface TCR/CD3 complexes with anti-CD3ε UCHT-1-scF_V_. Site-specific fluorophore conjugation of both constructs, guided by structural and functional considerations, yielded FRET yields that allowed unambiguous identification of single-molecule FRET signals with acceptable to high signal-to-noise ratios. Importantly, neither UCHT-1-scF_V_-CD3ε binding nor C-terminal fluorophore labeling of the HLA-associated β2m affected in our hands T-cell antigen sensitivities, which underscores the robustness of our experimental system.

FRET-based single-molecule analysis revealed that TCR:pMHC interactions with half-lives in the range of 1-2 seconds at 37°C supported the sensitive detection of a single antigen (RA 14 TCR-A2/CMW, 1E6-A2/RQW). In contrast, autorecognition (1E6 TCR-A2/PPI) relied on considerably shorter-lived interactions with half-lives around 300 milliseconds. Consistent with the kinetic proofreading model, short lifetimes were associated not only with low stimulatory potency but suggested furthermore thresholds in bond duration to be overcome for effective signaling.

In principle and consistent with TCR:pMHC catch-bond behavior, synaptic lifetimes for the RA14 TCR engaging A2/CMV exceeded in our hands at 37°C those measured by SPR by a factor of ∼2. However, factors other than forces may have contributed to the observed increase in bond duration within the synapse. Of note, binding partners lose at least one degree of freedom for bond-dissociation when embedded within opposed plasma membranes and the confines of the immunological synapse. Testing for catch-bond-dominated signaling-competent TCR-engagement will include (i) measuring the mechanical forces applied to individual A2/CMV-engaged synaptic RA14 TCRs and (ii) correlating them to bond lifetimes measured simultaneously. Using a FRET-based sensor, we have recently spatiotemporally recorded forces as they were exerted on individual TCRs within the immunological synapse of CD4+ murine T-cells ^12^. These forces amounted to a maximum of two piconewtons, contrasting the 15 piconewton force regimen reported previously for TCR:pMHC catch bonds to take effect ^21^. Also, recent evidence from *in vitro* experimentation argued in favor of accessory proteins shielding forces exerted on synaptic TCR:pMHC complexes rather than TCR-exerted forces being critical for sensitized antigen detection ^49^. More detailed studies will hence be required to reconcile the observed apparent differences in TCR:pMHC-bond stability measured *in vitro* and *in situ* and clarify the role of TCR-exerted forces on bond duration and signaling output.

Relevant in this context, CD8:MHCI engagement appeared as a *conditio sine qua non* for the single molecule sensitivities exhibited by RA14 T-cells facing A2/CMV and 1E6 T-cells scanning A2/RQW. In fact, disrupting A2:CD8 binding rendered antigen thresholds increased by up to three orders of magnitude. In contrast, inefficient autoantigen detection by 1E6 T-cells was only five-fold reduced, a perceived difference - at least in this study - which highlights the unique contribution of CD8-engagement not only to highly sensitized antigen recognition but also antigen discrimination. Based on the use of adhesion frequency assays, Zhu and colleagues proposed CD8-co-engagement as a critical contributor to catch-bonding between agonist pMHCs and their cognate TCRs ^37^. Contrary to this concept, not only the agonist (RA14 TCR:A2/CMV or 1E6 TCR:A2/RQW) but also the barely stimulatory autoreactive (1E6 TCR-A2/PPI) interaction benefited from CD8 co-engagement. While the extent to which catch-bonding contributed to the observed increase in synaptic bond lifetimes remains to be clarified (see above), our measurements challenge the previously proposed role of CD8-co-engagement in increasing solely the bond duration of agonist:TCR interactions ^37^.

Alternatively, simultaneous binding of TCR and CD8 to the same pMHC-target may stabilize the TCR:pMHC interaction for other yet undetermined reasons. When measured *in vitro*, CD8 affinity towards A2 amounted to ∼140µM ^46^, a value that exceeds the affinity determined via SPR for the 1E6 TCR-A2/PPI interaction (∼400 µM). As evidenced by the recruitment- and FRET-based analyses shown in this study, CD8-MHC binding supports even short-lived TCR:pMHC interactions and - most probably - vice versa. To our surprise, we failed to observe any synaptic accumulation of MHCI in the absence of TCR-matched antigen, suggesting that at least in the context of the immunological synapse CD8 does not interact with MHCI in an antigen-independent manner, or only very infrequently. The obvious contribution of CD8:MHCI-binding to the TCR:pMHCI-lifetime, as witnessed in this study, together with the apparent lack of CD8:MHCI-binding in the absence of antigen suggest an orchestrated mode of CD8-engagement in the presence of antigen. Definitive proof will however require direct monitoring of synaptic formation and dissociation of binary CD8-MHC and pMHC-TCR complexes and ternary CD8-pMHC-TCR complexes, ideally in a simultaneous manner, a technically ambitious endeavor outside the scope of this study.

Based on our results, we surmise that central tolerance mechanisms establish exceedingly high activation thresholds for self-antigens by effectively weeding out genetically recombined TCRs that mediate more sensitive autoantigen detection. In the context of the measurements presented, it will be informative to determine not only the copy number of individual endogenous pMHCs being displayed on the surface of mTECs but also the heterogeneity of AIRE-induced gene expression within the mTEC population. Such insights will help quantitate the effectiveness of central tolerance, in particular with knowledge of how frequently any given thymocyte will encounter mTECs displaying a given autoantigen. They will furthermore allow for estimates concerning the minimal number of autoantigens still tolerated by any scanning T-cell clone, ultimately defining the boundaries within which central tolerance operates. Notwithstanding, our measurements render it plausible that tissues featuring high copy numbers of individual pMHC-species, e.g. cells originating from protein- or peptide-secreting glands, are particularly susceptible to destruction by autoreactive T-cells, especially when the targeted autoantigen is expressed on mTECs at significantly lower numbers. A testable hypothesis relating to the clonality of autoreactive patient T-cells can be derived from (i) the high number of autoantigens required for T-cell activation and (ii) the considerable cross-reactivity of TCRs towards structurally unrelated foreign antigen ^50^. In essence, the rather low frequency of TCRs meeting these criteria suggests that the onset of many T-cell mediated autoimmune diseases may in fact be driven by only a few, if not even a single T-cell clone. If true, this could provide opportunities for identifying autoantigens based on high transcription levels in diseased organs and lower expression in mTECs. Once defined, autoreactive T-cell clones could be isolated based on pMHC-tetramer-binding, characterized and ultimately be therapeutically targeted in an antigen-or TCR-specific manner, as has been previously been reported with the use of T-cells modified with biomimetic five-module chimeric antigen receptors ^51^.

Irrespective of the molecular TCR-trigger mechanisms involved, our study lays bare the main difference separating sensitized from insensitive antigen recognition with regard to underlying membrane-proximal signaling. 1E6 T-cells required in a setting of autoreactivity at any time 100-times as many ligand-engaged TCRs as RA14 T-cells engaging A2/CMV. After compensating for differences in TCR-antigen affinities by means of experimentally adjusted TCR occupancies, synaptic CD3-ITAM phosphorylation levels were no longer distinguishable between the autoreactive and the anti-viral setting. In agreement with this, T-cell activation as measured via calcium signaling was equally affected in scenarios of viral and autoimmune recognition by partial Lck-inhibition. We therefore conclude that the average rate of antigen-triggered CD3 phosphorylation was similar, if not equal, when normalized to the total number of synaptic (i.e. frequency and duration) TCR:pMHC engagements. However, despite comparable pITAM-levels, resulting ZAP70 activation levels were at least two-fold lower in synapses of A2/PPI-engaged 1E6 T-cells. Of note, ZAP70 contains two SH2 domains, both of which must engage phospho-tyrosine residues within each ITAM for relatively stable ZAP70-pITAM association and further ZAP70 activation ^52^. Short-lived TCR interactions with autoantigens hence generate most likely numerous incompletely phosphorylated TCR/CD3 complexes, which fail to stably recruit ZAP70 and which remain, as a consequence, signaling-incompetent. Conversely, more stimulatory antigens trigger by means of longer-lasting TCR interactions fully phosphorylated and signaling-competent TCR/CD3 complexes, even in low antigen abundance. Since we could not detect pITAM- or pZAP70-immunofluorescence above background levels at agonist densities at which RA14 T-cells already showed a robust calcium response, we consider it highly plausible that the generation of only a small number of signaling-competent (i.e. fully phosphorylated) TCR/CD3 complexes supports uninterrupted T-cell activation. To test this hypothesis and advance rational approaches for T-cell-based immunotherapies, it will be critical to quantify the exact number of fully triggered TCR/CD3 complexes required for T-cell activation. Predictive parameters –to be assessed as a function of antigen quality and quantity – will likely include (i) the time required to complete the phosphorylation of individual TCR/CD3 complexes in the presence and absence of CD8-binding, (ii) their lifetime after antigen-dissociation, as well as (iii) degrees of heterogeneity in ITAM-phosphorylation.

In summary, quantitative and advanced molecular imaging has allowed us to compare T-cell antigen recognition scenarios at the level of synaptic TCR:pMHC binding and membrane proximal signaling in settings of anti-viral immune responses and autoimmunity. Our study points to pivots that appear relevant to establishing exquisite T-cell antigen sensitivity while simultaneously quenching autoreactivity: (i) the large dynamic range of TCR:pMHC engagement kinetics supporting productive receptor triggering, (ii) the role of CD8-co-engagement in discriminating antigenic from endogenous pMHCs, and (iii) the dynamics underlying pITAM-dependent ZAP70-activation as a critical checkpoint in T-cell activation. Gained insights further a quantitative understanding of T-cell antigen recognition dynamics which support ultra-sensitive antigen detection in the context of TCR-cross-reactivity and central tolerance, with direct implications for T-cell function in settings of infection, autoimmunity, cancer, organ transplantation and therapeutic T-cell engineering.

## METHODS

### Lead contact

Further information and requests for resources and reagents should be directed to and will be fulfilled by the lead contacts, Janett Göhring and Johannes Huppa.

### Materials availability

Reagents generated in this study will be available upon request. DNA sequences have been deposited at GenBank and accession numbers are listed in the key resources table.

### Data and code availability

- Raw microscopy data have been deposited at Figshare and are publicly available as of the date of publication. Flow cytometry data (FCS files) have been deposited at Flow repository and PDB files have been deposited at PDB. All accession numbers are listed in the key resources tables.
- This paper does not report original code.
- Any additional information required to reanalyze the data reported in this paper is available from the lead contacts upon request.

## EXPERIMENTAL MODEL AND STUDY PARTICIPANT DETAILS

### Primary cultures and cell lines

We employed CRISPR-Cas9 orthotopic TCR exchange in primary human T-cells to generate CMV-specific RA14 and autoreactive, preproinsulin (PPI)-specific 1E6 T-cells. Peripheral blood samples were obtained from blood donations from anonymous donors providing informed consent according to our Ethics Guidelines (Ethics Committee of the Medical University of Vienna, EK Nr: 2001/2018). All cell lines and primary cells were cultured at 37°C, 5% CO_2_ and 95% relative humidity in a tissue culture incubator. We cannot report the sex of our primary cells as blood donors remain anonymous in accordance with our ethics guideline.

### Microbial culture conditions

Escherichia coli (E. coli) cultures were grown at room temperature, shaking with 200 rpm. Insect cells were cultured at 26-28°C in a non-humidified incubator without CO_2_ control, shaking with 130 rpm.

### Recombinant proteins

Recombinant proteins were produced in E. coli-BL21 (DE3) or High Five insect cells, as indicated in the methods section.

## METHOD DETAILS

### Protein production and purification

HLA.A*0201 (A2) proteins for pMHC-tetramer staining or SLB functionalization were produced via bacterial expression in a pET-28b vector containing the cDNA encoding the extracellular domain of A2 (UniProt: P01892) without a leader sequence and followed by a C-terminal Avi-tag (GLNDIFEAQKIEWHE) for site-specific biotinylation or 12xHis-tag for SLB-incorporation, respectively. For A2^ΔCD8^ constructs, we mutated the CD8 binding site on the A2 heavy chain by inserting two mutations in position 227 and 228 (D227K and T228A). The cDNA encoding β2m (UniProt: P61769) without a leader sequence was cloned for bacterial expression in a pHN1 vector. Recombinant pMHCs were expressed as inclusion bodies in Escherichia coli. After a series of clean-up steps and subsequent resuspension in 6M guanidium hydrochloride, pMHC were refolded in presence of the corresponding peptides, pp65-CMV (NLVPMVATV), preproinsulin 15-24 (PPI_15-24_; ALWGPDPAAA) or the preproinsulin 15-24 altered peptide variant RQW (RQW: RQWGPDPAAV) to form A*0201-peptide complexes as described ^53,54^. After completion of the refolding reaction, the solution (330 ml) was dialyzed against 20 l of PBS. For site-specific biotinylation, PBS was exchanged against 50 mM bicine buffer (pH 8.3) supplemented with 200 mM potassium glutamate. The protein solution was concentrated to 40 µM to ensure efficient biotinylation (according to reaction conditions for Bir A500 by Avidity LCC). Next, Biomix B (10x stock: 100 mM ATP, 100 mM MgOAc and 500 µM biotin) and Glutathione -S-transferase (GST)-tagged BirA enzyme (2.5 µg per 10 nmol protein solution) were added to the protein solution and incubated for 60 minutes at 30°C. GST-tagged BirA was removed by incubation with glutathione agarose (Thermo Scientific) and by subsequent 0.22 µm filtration. 12x-His-tagged A2 complexes were directly purified via affinity chromatography using a Ni^2+^-NTA agarose column after dialysis. To exclude protein aggregates or protein dimers, affinity-purified protein was subjected to Superdex 200 (Superdex 200, 10/300 GL, Cytiva) size-exclusion chromatography (SEC). Protein purity and successful biotinylation was verified via SDS-PAGE and a streptavidin gel shift assay followed by silver or colloidal Coomassie blue staining.

Recombinant UCHT-1 scF_V_ was expressed as inclusion bodies in a pET-21a+ vector in E. coli-BL21 (DE3). The refolding reaction was performed as described in Tsumotot et al ^55^. In brief, a slow buffer exchange is performed over the course of 6 days, resulting in a stepwise removal of denaturing reagents and addition of oxidizing agents, concluding with a dialysis step in PBS. Refolded UCHT-1 scF_V_ was further purified by SEC.

ICAM-1-12xHis and B7-1-12xHis for SLB functionalization were produced in High Five insect cells (BTI-Tn-5B1-4, Thermo Fisher Scientific) infected with viral particles of the Baculovirus Expression System as described ^56^.

### Fluorophore conjugation

A2 complexes used for microscopy were fluorescently labeled either via amine-reactive N-hydroxysuccinimide ester or site-specifically fluorescence-tagged, whenever required, via sulfhydryl-reactive maleimide chemistry.

For N-hydroxysuccinimide ester labeling, protein aliquots of 100-300µg in PBS were adjusted to a pH of 8.3 by adding freshly prepared NaHCO_3_ (0.1 M final concentration) and incubated for 20 minutes at RT with Alexa Fluor 647-NHS dye (Thermo Fisher Scientific) in six-fold molar excess.

For site-specific maleimide labeling, protein aliquots of 100-300 µg in PBS were mixed with freshly dissolved TCEP in HEPES buffer pH 7.0 (50 µM final concentration) and Alexa Fluor 647-maleimide dye in ten-fold molar excess (Thermo Fisher Scientific) for 2 hours at RT. Labeled proteins were separated from free dye solution by Superdex 75 (Superdex 75, 10/300 GL, Cytiva) size-exclusion chromatography. The degree of dye-conjugation was determined by photo-spectrometry. Labeled proteins were stored in PBS supplemented with 50% glycerol at - 20°C until further use.

### Generation of monoclonal T-cells via CRISPR-Cas9 mediated TCR exchange

Frozen PBMCs from A2-negative donors were thawed, rested in RPMI with 10% human serum and 50 U/ml human IL-2 overnight and enriched for CD8+ T-cells via the Magnisort^TM^ human CD8+ T-cell enrichment kit (Thermo Scientific). CD8 T-cells were stimulated with plate-bound OKT3 antibody (5 µg/ml) and soluble anti-CD28 antibody (2 µg/ml) in the presence of 300 U / ml human IL-2 and 5 ng/ml IL-15 in RPMI supplemented with 10% human serum for 48 hours prior to electroporation. For targeted TCR alpha- and beta-chain double knock-out, ribonucleoprotein (RNP) complexes were produced using the following CRISPR RNAs (crRNA): 5′-GGAGAATGACGAGTGGACCC-3′ for TRBC (targeting both TRBC1 and TRBC2) and 5′-AGAGTCTCTCAGCTGGTACA-3′ for TRAC ^57^. crRNAs (80 µM, Integrated DNA Technologies) were incubated with trans-activating RNA (tracrRNA, 80 µM, Integrated DNA Technologies) at 95°C for 5 minutes and subsequently allowed to cool down to RT. gRNAs were gently mixed with High-Fidelity Cas 9 (24 µM, Integrated DNA Technologies) and incubated at RT for 20 minutes to yield RNP complexes with 12 µM Cas9, 20 µM gRNA and 20 µM electroporation enhancer (Integrated DNA Technologies). For CRISPR-Cas9 mediated TCR knock-in via homology-directed repair (HDR) double-stranded DNA, PCR products with homology sequences located 5′ and 3′ of the introduced TCR, which targeted the first exon of the endogenous TRAC locus were produced as described ^27^. Double-stranded DNA templates were designed according to the following structure: left homology arm, P2A site, TCR beta chain, T2A site, TCR alpha chain, Stop codon, Poly A site, right homology arm. Double-stranded DNA templates were amplified by PCR and purified with the use of a Wizard® SV Gel and PCR Clean-Up kit (Promega). CD8+ T-cells were electroporated using the Amaxa Nucelofector II (program T23) in 100 µl electroporation buffer per 5 × 10^6^ T-cells (Human T-cell nucleofection Kit, Lonza) in the presence of 2 µg double-stranded DNA template and Cas9-RNP complexes. After electroporation, cells were cultured in RPMI supplemented with 10% human serum and 180 U / ml human IL-2 prior to flow cytometry analysis and subsequent FACS sorting.

### Tetramer-based staining of T-cells

pMHC-tetramers were formed by mixing biotinylated pMHC monomers (produced in-house as described above) in a 4:1 molar ratio with Streptavidin-PE or Streptavidin-APC (both BioLegend) on ice. Streptavidin was added to the pMHC monomers in 10 portions every 10 minutes. To remove incompletely oligomerized complexes, protein solutions were incubated with biotin-agarose (Thermo Fisher Scientific) and purified by spin-filter centrifugation. T-cells were incubated with 20 µg/ml tetramers on ice in PBS containing 1% bovine serum albumin (BSA) for 45 minutes before washing and subsequent flow cytometric analysis.

### Fluorescence activated cell sorting (FACS)-based enrichment of RA14 and 1E6 T-cells

Five days after CRISPR-Cas9-mediated TCR replacement, T-cells were stained with PE- and APC-conjugated pMHC-tetramers at a concentration of 0.5µg tetramer per 10^6^ T-cells on ice in PBS containing 1% bovine serum albumin (BSA) and 200 mM ethylenediaminetetraacetic acid (EDTA) for ten minutes prior to the addition of Alexa Fluor 488 labeled anti-human αβTCR-reactive IP26 mAb (Biolegend, final concentration: 20 µg/ml) and APC-eFluor780 labeled CD8α-reactive OKT8 mAb (eBioscience, final concentration: 2.5 µg/ml) antibodies. RA14 and 1E6 T-cells were sorted as tetramer-specific CD8- and TCR-positive cell populations.

### Antigen-specific expansion of RA14 and 1E6 T-cells

For peptide-specific expansion, we co-cultured TCR-exchanged T-cells with the human lymphoblastoma cell line K562 transduced to express HLA-A*0201 (kind gift of Peter Steinberger, Medical University of Vienna ^58^) pulsed with 10 µM of the CMV pp65 or the PPI15-24 peptide for 2 hours at 37°C and irradiated with 80 Gray. K562 cells were added to T-cells at a ratio of 3:1 in RPMI supplemented with 10% human serum and 180 U/ml human IL-2 (Proleukin, Norvatis), and expansion was completed after 7-10 days.

### Preparation of SLB

SLBs were prepared and functionalized with proteins of interest as described ^59^. Briefly, 98 mol-% POPC and 2 mol-% Ni-NTA-DGS, both dissolved in chloroform, were mixed and then dried under vacuum in a desiccator overnight. The lipid mixture was resuspended in degassed PBS and sonicated under nitrogen in a water bath sonicator (Q700; QSonica) for 45min to form small unilamellar vesicles (SUV). To remove non unilamellar vesicles, the solution was centrifuged for 1h at 37000rpm at RT and subsequently 8 hours at 43,000 rpm at 4 °C using a Sorvall RC M150GX ultracentrifuge with a S150AT-0121 rotor (Thermo Fisher Scientific). The supernatant was 0.2 µm filtered and stored under nitrogen at 4 °C for up to six months.

For bilayer preparation, glass slides were plasma cleaned (Zepto plasma cleaner, Diener electronics) for 20min, and glued to the bottom of 8-well or 16-well Nunc Lab-Tek chambers (Thermo Fisher Scientific) using Picodent twinsil (Picodent). The SUV solution was diluted 1:20 in PBS, added to each well and allowed to form a contiguous SLB within 15 minutes at RT. Chambers were washed with 20ml PBS to remove the excess of lipid vesicles, before 12x-His-tagged proteins of interest were added to the well and incubated for 45 minutes at RT in the dark. To remove unbound protein, chambers were rinsed with 20ml PBS and kept in the dark until usage within 6h. Protein density on the SLB was determined prior to every experiment by dividing bulk florescence intensity by single-molecule florescence intensity. For cell imaging PBS was changed to Hank’s Balanced Salt Solution (HBSS) supplemented with 2 mM of CaCl_2_, 2 mM of MgCl_2_ and 2% FCS.

### Microscopy setup

Microscopy was performed with the use of two custom-built microscope setups for TIRF-based imaging or calcium imaging.

Setup #1: For TIRF-based imaging, SLB were illuminated in TIRF mode with an inverted microscope (Eclipse Ti-E, Nikon Instruments) equipped with a chromatically corrected 100x TIRF objective (CFI SR Apo TIRF 100× Oil NA:1.49, Nikon Instruments) and diode lasers featuring the wavelengths of 488nm, 514nm, 642nm (iBeam smart Toptica) or 532nm and 561nm (OBIS) for excitation as well as a xenon lamp (Lambda LS lamp, Sutter Instruments). Laser excitation was cleaned up by clean-up filters (Chroma) matching the individual laser wavelengths and individual laser lines. Lamp excitation was filtered via the following excitation band pass filters: 340/26, 387/11, 370/36, 474/27, 554/23 and 635/18 (all Chroma). Laser and lamp excitation light was separated by beam splitters zt405/488/561/647rpc, zt405/488/532/640rpc and zt405/514/635rpc (Chroma). For FRET-based measurements, the fluorescence emission was split into two spectral channels by an Opto-split 2 (Cairn Research) equipped with a beam splitter (ZT640rdc, Chroma) and appropriate emission bandpass filters (ET575/50 or ET640SP, ET655LP, Chroma). Images were recorded using an Andor iXon Ultra 897 EMCCD camera (Oxford Instruments). Image acquisition was controlled with the use of an 8-channel DAQ-board PCI-DDA08/16 (National Instruments) and the microscopy automation and image analysis software MetaMorph (Molecular Devices) to program and apply timing protocols and control all hardware components.

Setup #2: For calcium recordings, a second inverted microscope (DMI4000, Leica Microsystems) with a chromatically corrected 20x objective (HC PL FLUOTAR 20×/0.50 PH2∞/0.17/D, Leica Microsystems) and an EL6000 mercury lamp (Leica Microsystems) were used. The microscope was furthermore equipped with a fast filter wheel containing 340/26 and 387/11 excitation bandpass filters (both Leica Microsystems) and the beam splitters calcium cube FU2 (Leica Microsystems) and ZT405/488/532/647rpc (Chroma). Images were recorded with a Prime 95B sCMOS camera (Photometrics). The operation of hardware components was controlled via µManager ^60^.

Temperature control was conducted using a closed stage-housing system from Pecon (setup #1) and Leica (setup #2).

### Determination of antigen density

Quantification of antigen densities on the SLB was performed as described ^59^. Briefly, bulk fluorescence intensity was background-corrected and divided by single molecule fluorescence intensity. To measure bulk fluorescence intensity, a region of interest (ROI) of 100×100 pixels was placed over the maximal laser illumination spot and intensity counts were integrated as well as subtracted for background intensity, which was recorded under identical imaging conditions on antigen-free SLBs.

For determination of single molecule intensity, SLBs with antigen densities which allow for visualization of separate diffraction-limited single molecule events with non-overlapping point-spread functions were used. Again, a ROI of 100×100 pixels was placed over the maximal laser illumination spot and further analyzed by the ImageJ Plugin ThunderStorm ^61^, which enables the localization of individual single molecule events. The fluorescence intensity of all detected single molecule events was next plotted as a histogram, and the mean fluorescence intensity was calculated. Recorded bulk fluorescence intensities were divided by the mean fluorescence intensity values derived from recorded single molecule events (n>100) to yield the number of antigens per ROI, which was then divided by the pixel area multiplied by the camera-specific pixel to µm^2^ conversion factor (1 µm^2^ ≙ 6.25 x 6.25 camera pixels) to arrive at the antigen density (molecules µm^-2^).

### Determination of protein mobility by FRAP

Fluorescence recovery after photobleaching (FRAP) experiments were performed as described^59^. Briefly, a circular aperture was placed into the laser beam path and x/y-positioned to the spot of maximum laser illumination. Three images were recorded at laser intensities that keep fluorophore bleaching at a minimum, followed by a bleach pulse with maximum laser intensity to ablate fluorescence quantitatively and a consecutive fluorescence image to confirm successful fluorophore bleaching, before fluorescence-labeled proteins were allowed to diffuse back into the bleached area. Fluorescence recovery was monitored for 10 minutes at 1-minute time intervals. Measured intensities were background-corrected and normalized against the mean of the three intensities recorded prior to fluorophore ablation.

### Calcium imaging

Intracellular calcium levels were measured with the use of the ratiometric calcium sensitive dye Fura-2-AM as described in ^62^. Briefly, a total of 1 × 10^6^ T-cells were stained in 1 ml PBS supplemented with 5 µM Fura-2-AM (Thermo Fisher Scientific) for 20 minutes at 37°C and washed once in 10 ml ice-cold imaging buffer, which consisted of Hank’s Balanced Salt Solution (HBSS) supplemented with 2 mM of CaCl_2_, 2 mM of MgCl_2_ and 2% FCS. SLBs were immersed in imaging buffer and allowed to prewarm to 37°C, directly before approximately 100,000 T-cells per well were spotted onto the SLB. By employing an automated XY stage (Leica Microsystems), time-lapse images of the calcium response of T-cells upon SLB contact were recorded at up to six different XY stage positions per run with alternating 340 and 387 nm excitation every 15 seconds for 20 min, using an ultraviolet-transmissive 20× objective (HC PL FLUOTAR 20×/0.50 PH2∞/0.17/D, Leica Microsystems).

For calcium analysis, we tracked individual cells using a published particle tracking algorithm^31^. Tracking parameters were chosen so that only T-cells that had been in contact with the SLB for at least 12 minutes were included in the analysis, which resulted in an average of 250 (ranging from 80 to 550) tracks for each experimental condition chosen. For each track chosen we determined the mean Fura-2 ratio value in close vicinity of the peak of the calcium response (median of peak + 10 frames, corresponding to a time interval of 2.5 minutes). Population analysis of the generated ratiometric images was conducted with the use of a custom-built MATLAB software script based on the previously described principles ^63^. Briefly, ratiometric images were normalized to the population median of the negative control, which consisted of an antigen-free SLB decorated only with ICAM-1 and B7-1, and which was recorded under identical imaging conditions. The individual trajectories of the population were synchronized to the first camera detection of the cell on the SLB. In this fashion, we classified T-cells, which were found for 80% of their trajectory above a defined activation threshold (determined with the use of the negative control as reference and in most cases defined as Fura-2 ratio of >1.3), as activated. The activation threshold is determined as the maximum difference between the negative and positive control by means of receiver-operator characteristics (ROC) analysis for each experiment. The maximum allowed false positive (FP) rate is 0.05, however, in a routine experiment the activation threshold would be set to yield even lower FP rates.

To assess the quality of T-cell activation, a histogram of the mean Fura-2 ratio for each cell was generated (over the range of 10 frames starting from the maximum Fura-2 ratio of each individual cellular trajectory). To ensure only activated cells were included in the calculation, a new histogram was generated using the activation threshold determined by previous population analysis (ROC cut-off at 1.3). The mean Fura-2 value *μ* of the activated population was estimated based on the mean of the histogram (*μ* = (∑ *m*_*i*_*n*_*i*_)⁄*N*, where *m* and *n* is the midpoint and frequency of the *i*^th^ bin of the histogram, respectively, and *N* is the total sample size).

### Quantitation of IFN-γ secretion after antigenic stimulation of T-cells on SLBs

Prior to T-cell seeding, antigen densities on SLBs were determined as described above. 100.000 T-cells were spotted per well on SLBs equipped with various antigen densities of A2 with the corresponding peptides, ICAM- and B7-1, and incubated at 37°C for 30 min. After this first incubation of the cells in 100 µl imaging buffer (HBSS supplemented with 2 mM of CaCl_2_, 2 mM of MgCl_2_ and 2% FCS), the wells were filled with additional 300 µl human T-cell medium (RPMI 1640 medium supplemented with 25 mM of HEPES, 10% human serum, 100 U ml−1 of penicillin/streptomycin, 2 mM of L-glutamine and 50 µM of 2-mercaptoethanol) and incubated for 72 hours at 37°C. Then, the supernatants were collected and stored at -20°C until further use. For determination of secreted Interferon gamma (IFNγ), 100µl of the collected supernatants were analyzed via ELISA (ELISA MAX Deluxe Set for human IFNγ, BioLegend).

### Killing assay of peptide-pulsed K562 cells co-cultured with RA14 and 1E6 T-cells

K562 cells expressing HLA.A2 and luciferase firefly-BFP was pulsed with different concentrations (1 µM to 0.001 nM in 1:10 dilutions) of CMV pp65, PPI_15-24_ or the altered peptide ligand RQW for 1 hour at 37°C in RPMI 1640 medium supplemented with 10% human serum, 100 U ml^−1^ penicillin/streptomycin, 50 µM of 2-mercaptoethanol (Thermo Fisher Scientific). After peptide-pulsing 75 µg ml^−1^ D-luciferin firefly (Biosynth) was added. To assess their target cell killing performance, 30.000 RA14 or 1E6 T-cells were added to peptide-pulsed K562 cells at an effector:target ratio of 1:1 in triplicates. As the maximum killing control, K562 cultured in 50% DMSO were used. Spontaneous cell death was assessed in wells with K562s only (duplicates). K562 viability was measured by relative light units (RLU) after 6, 8 and 19 hours on the microplate reader (Mithras LB 940). Specific cell lysis was determined using equation (1):

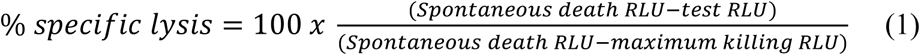

### Quantification of T-cell antigen recruitment

The number of recruited antigens per synapse of T-cells encountering different antigen densities on SLBs was determined by fluorescence quantification after background correction. 15.000 RA14 or 1E6 T-cells per condition were spotted onto SLBs and allowed to adhere for 3 min.

IRM images were acquired to determine the synapse area (cell ROI) and recruitment of Alexa Fluor 647-labeled A2 molecules was assessed by recording a TIRF image using 640nm laser illumination at a power density of 0.1 kW cm^-2^ and 20 ms sample exposure time (ET700/75 bandpass filter). For quantification of ligand recruitment, intensity counts of cell ROIs were integrated and divided by single-molecule intensity to arrive at the number of molecules within the cell ROI. Quantification of a cell-free area next to the cell ROI served as negative control for background subtraction. To compensate for the Gaussian laser profile and resulting lower illumination power densities outside the cell ROI, we employed flatfield correction. When working with SLB antigen densities too high for separating diffraction-limited single events, we applied the following formula for Flatfield correction (equation 2):

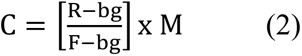

with *C*= corrected image, *R* = raw image before correction, *F* = flatfield image (resulting from the median of at least 10 images exhibiting homogenous intensity distributions), *bg* = camera background, *M* = average intensity of flatfield image (*F*) – camera background (*bg*) . The number of ligands next to the cell ROI (background) was then subtracted from the number of ligands within the cell ROI to arrive at the number of recruited antigens per T-cell synapse. For lower antigen densities on SLBs, individual molecules within the cell ROI and next to the cell ROI (background) were quantified by ThunderStorm tracking (see above).

Fold increase values of initial SLB antigen densities were calculated by correcting the number of molecules per synapse / next to the synapse for camera pixel size to arrive at the number of molecules µm^-2^. Recruited molecules within the cell ROI were divided by background values to obtain fold increase ratios. The number of recruited antigens as well as fold increase values were plotted against antigen densities on SLBs and error bars display the s.e.m.

### Quantitation of TCR/CD3 complex decoration by UCHT-1 scF_V_

RA14 or 1E6 TCR T-cells were stained with increasing concentrations of Alexa Fluor 647-labeled UCHT-1 scF_V_ or Alexa Fluor 647-labeled anti-human TCR α/β antibody (clone IP26, BioLegend). The mean fluorescent intensity of stained T-cells was measured by flow cytometry. MFI values were converted to molecules of equivalent soluble fluorochrome (MESF) units with the use of simultaneously measured quantitation beads (Quantum MESF Alexa Fluor 647; Bangs laboratories).

### Ensemble FRET imaging

For ensemble FRET measurements, T-cells were stained with site-specifically Alexa Fluor 647-labeled UCHT-1 scF_V_ for 30 minutes on ice, washed twice with imaging buffer (HBSS supplemented with 2 mM of CaCl_2_, 2 mM of MgCl_2_ and 2% FCS), and spotted onto SLBs that had been equipped with Alexa Fluor 647-labeled antigen (A2* with the corresponding peptides), ICAM-1 and B7-1. After allowing the T-cells to adhere to the SLB for 2 minutes synaptic FRET was recorded for up to 15 minutes. FRET efficiencies between Alexa Fluor 555-labeled UCHT-1 scF_V_ decorated T-cells (donor) and bilayer-resident A2*/peptide-Alexa Fluor 647 (acceptor) were determined by Donor Recovery After Acceptor Photobleaching (DRAAP). In principle, the donor channel as well as the acceptor channel were recorded prior to and post acceptor photobleaching. All exposures were conducted in TIRF mode using 50 ms exposure times for the donor and acceptor channel and a bleach pulse of 200 ms.

For FRET efficiency determination, a custom-built MATLAB software was used that is based on described principles ^64^. Briefly, the donor intensity before acceptor photobleaching was subtracted from the donor intensity after acceptor photobleaching. The resulting difference was then divided by the donor intensity after acceptor photobleaching to obtain the FRET efficiency. In addition, the pixel-averaged fluorescent intensity of the donor signal of each cell was corrected for photobleaching, cross-excitation as well as bleed-through based on ^64^.

In a last step, FRET efficiencies were corrected for the degree of labeling of engaging antigens, as unlabeled ligands engage TCRs but do not result in FRET.

### smFRET imaging and determination of dissociation rates from smFRET

To limit bleed-through from the donor fluorophore into the FRET channel for the detection of single molecule FRET events, UCHT-1 scF_V_-TCR fluorescence decoration was limited to 30% of all antibody-accessible TCRs on the T-cells by staining with a premixed cocktail of unlabeled and site-specifically Alexa Fluor 555-labeled UCHT-1 scF_V_ at a ratio of 2:1.

SLBs were prepared to feature fluorescent antigen densities of approximately 10 molecules µm^-2^. To ensure activating conditions, unlabeled ligands (A2* with the according peptides) were incorporated into the SLB to give rise to stimulatory antigen densities of >100 molecules µm^-2^. For single-molecule FRET measurements, T-cells decorated with site-specifically labeled UCHT-1 scF_V_ (as described above) were spotted on SLBs with a fluorescent antigen density of approximately 10 molecules µm^-2^. To ensure activating conditions, unlabeled ligands (A2* with the according peptides) were added to provide stimulatory densities of >100 molecules µm^-2^ on SLBs.

For the determination of dissociation rates, typically 125 to 500 images of the FRET channel were recorded with 532nm excitation. The illumination power density was set to 0.1 kW/cm^2^ for 20 ms camera exposure times using different delay times (56 ms, 84 ms, 252 ms, 476 ms, 924 ms and 1820 ms). For especially short-lived interactions (A2^ΔCD8^/PPI) we increased the illumination power density to 2.5 kW/cm^2^ and reduced exposure times to 1ms employing different delay times (7 ms, 10 ms, 14 ms and 28 ms). Single-molecule FRET events had to appear and disappear in single, discrete steps to be classified as a true single-molecule event. Cells were imaged at 26°C and 37°C for no longer than 20 minutes after spotting. For analysis, our previously published analysis software based on the std and SciPy Python packages was employed ^45^. In brief, apparent (photobleaching-affected) FRET event half-lives were calculated for each delay time from single molecule track lengths using survival analysis. Physical half-lives and error estimates were obtained by fitting an appropriate model to the delay time vs. apparent half-live graph using a weighted non-linear least squares method.

### Quantitation of synapse-associated CD3ζ ITAM- and ZAP70-phosphorylation

Prior to T-cell seeding, antigen densities on SLBs were determined as described above. T-cells were spotted on SLBs equipped with various antigen densities of A2 with the corresponding peptides, ICAM- and B7-1, and allowed so settle and respond to the stimulus at 37°C for 10 minutes. Then, cells were fixed and permeabilized in one step by incubating SLBs for 20 minutes at RT with PBS supplemented with 4% formaldehyde, 1 mM of Na_3_O_4_V, 50 mM of NaF and 0.1% Triton X-100 Surfact-Amps (final concentrations; all from Thermo Fisher Scientific). After SLBs were rinsed with washing buffer, consisting of PBS supplemented with 3% bovine serum albumin (BSA), 1 mM of Na_3_O_4_V and 50 mM of NaF, cells were incubated overnight at 4 °C with unlabeled primary monoclonal antibodies reactive to rabbit anti-CD3ζ (phospho Y83) (clone EP776(2)Y; Abcam) plus mouse anti-phospho-CD3ζ (pY142) (clone K25-407.69; BD Biosciences) or mouse anti-phospho-CD3ζ (Tyr111, Tyr123) (clone EM-55; Thermo Fisher Scientific) plus rabbit anti-ZAP-70 phosphorylated at position 319 (clone 1503310; BioLegend). The next day, unbound antibody was removed by rinsing the sample with washing buffer. Secondary fluorescently labeled antibodies, goat anti-rabbit IgG (H+L) Alexa Fluor 647 (catalog no. A27040, Thermo Fisher Scientific) and goat anti-mouse IgG (H+L) Alexa Fluor 488 (catalog no. A28175, Thermo Fisher Scientific), were added and incubated for 3h at 4°C. After a final washing step, synapse associated phosphorylated CD3ζ-ITAM and ZAP-70 molecules were quantified by TIRF-based microscopy. The integrated intensity of the Alexa Fluor 647- or Alexa Fluor 488-conjugated secondary antibodies, detecting indicated phosphorylated CD3ζ ITAM and ZAP-70 molecules, were plotted against antigen density on the SLBs.

## STATISTICAL ANALYSIS

All statistical analyses were performed in GraphPad Prism or Python. Error bars in figures represent either standard deviation (SDev) or standard error of the mean (SEM) as indicated in the figure legends. TCR:pMHC binding half-lives were derived by MLE and fitted using a non-linear least squares regression analysis.

## ACKNOWLEDGMENTS

We kindly thank Peter Steinberger and his group for providing the K562 target cell line.

## AUTHOR CONTRIBUTIONS

Conceptualization: AP, VM, JG, JBH

Methodology: AP, VM, AR, RP, IDP, TP, CM, AR, YW, KS, DHB, JG, JBH

Investigation: AP, VM, AR, PF, CM, TP

Visualization: AP, VM, JG, JBH

Funding acquisition: YW, HS, LD, DHB, JG, JBH

Project administration: JBH

Supervision: CM, YW, DHB, KS, JG, JBH

Writing – original draft: AP, VM, JG, JBH

Writing – review & editing: AR, RP, YW, LD, CM, HS, DHB, KS, JG, JBH

## SUPPLEMENTARY FIGURE LEGENDS

**Figure S1.**
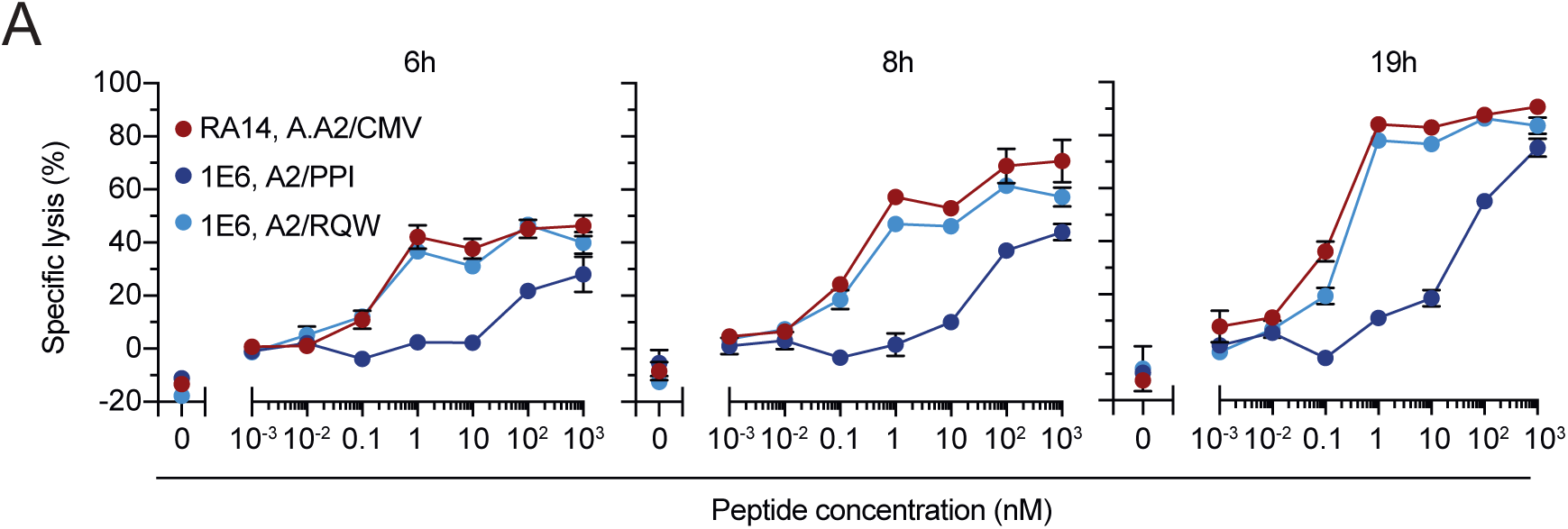
Assessment of killing capacity of RA14 and 1E6 T-cells towards antigen-pulsed target cell line. (**A**) Quantification of cytolytic activity of RA14 and 1E6 T-cells after stimulation with peptide-pulsed K562-HLA.A2 luciferase-expressing target cells in a 1:1 ratio for indicated times (6, 8 and 19 hours). Data shown of one experiment.

**Figure S2.**
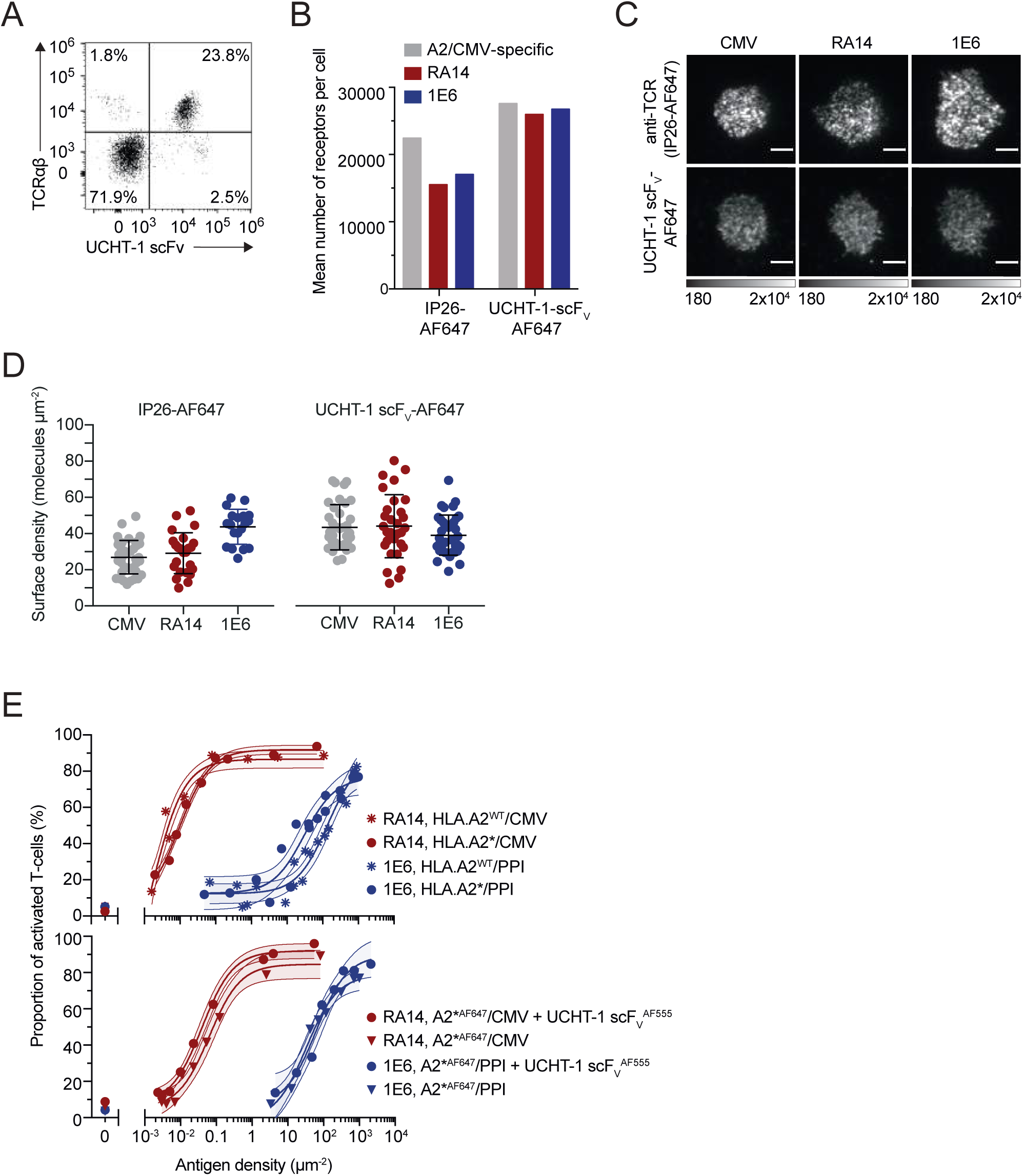
UCHT-1 scF_V_ specifically stains T-cells co-stained with an TCR-reactive antibody and does not alter antigen sensitivity of decorated T-cells. (**A**) Specificity of UCHT-1 staining is confirmed by co-staining of PBMCs with 40µg/ml UCHT-1 scFV-AF555 and 80µg/ml AF647-labeled TCRαβ reactive IP26 antibody. (**B**) Polyclonal A2/CMV-specific T-cells (grey) isolated from PBMCs via A2/CMV-tetramers, RA14 TCR (red) and 1E6 TCR (blue) T-cells were labeled with 80µg/ml TCR-reactive antibody IP26-AF647 (**left panel**) and 40µg/ml CD3-specific UCHT-1 scF_V_ AF647 (**right panel**) under saturating conditions. Staining concentrations have been determined by prior antibody titrations. Number of molecules per cell (MESF) were derived from calibration curves of AF647 intensity values using Quantum^TM^ AF647 MESF beads (Bangs Laboratories). (**C**) Polyclonal CMV-specific T-cells (CMV), which had been isolated from PBMCs via A2/CMV-tetramers, as well as RA14 and 1E6 T-cells were decorated with UCHT-1 scF_V_ or anti-TCR mAb (clone IP26) under saturating conditions, next allowed to spread on ICAM-1-functionalized SLBs and finally fluorescence intensity acquired in TIRF mode. Scale bar, 5µm. (**D**) TCR/CD3 surface densities were determined for polyclonal A2/CMV-specific (CMV, grey) as well as RA14 (red) and 1E6 (blue) T-cells with the use of fluorescence-tagged UCHT-1 scF_V_, anti-TCR mAb (clone IP26) and TIRF imaging as depicted in (C). (n ≥ 20 cells per condition). (**E**) Upper panel: Antigen sensitivities of RA14 (red) and 1E6 (blue) T-cells, as monitored via calcium imaging, were not affected by the use of site-specifically AF647-labeled A2/peptide complexes. Randomly labeled A2/peptide complexes (A2*) or a β2m-modified version of A2/peptide featuring a site-specific maleimide-based fluorochrome conjugation at an unpaired cysteine residue in the β2m subunit at position I2C (b2mI2C) (A2^wt^) were placed at indicated densities together with ICAM-1 and B7-1 on SLBs for stimulation of indicated T-cell populations which had been seeded on top. Lower panel: T-cell decoration with of UCHT-1 scF_V_ did not affect T-cell antigen sensitivity as monitored via calcium imaging of RA14 (red) and 1E6 (blue) T-cells confronted with SLBs featuring in addition to ICAM-1 and B7-1 the respective antigen at indicated densities. T-cells were first labeled with UCHT-1 scF_V_ (circles) or mock-stained (triangles) and then allowed to engage SLBs. T-cells confronted with antigen-free SLBs functionalized with ICAM-1 and B7-1 only served as negative control. Data shown of n= 1-2 experiments.

**Figure S3.**
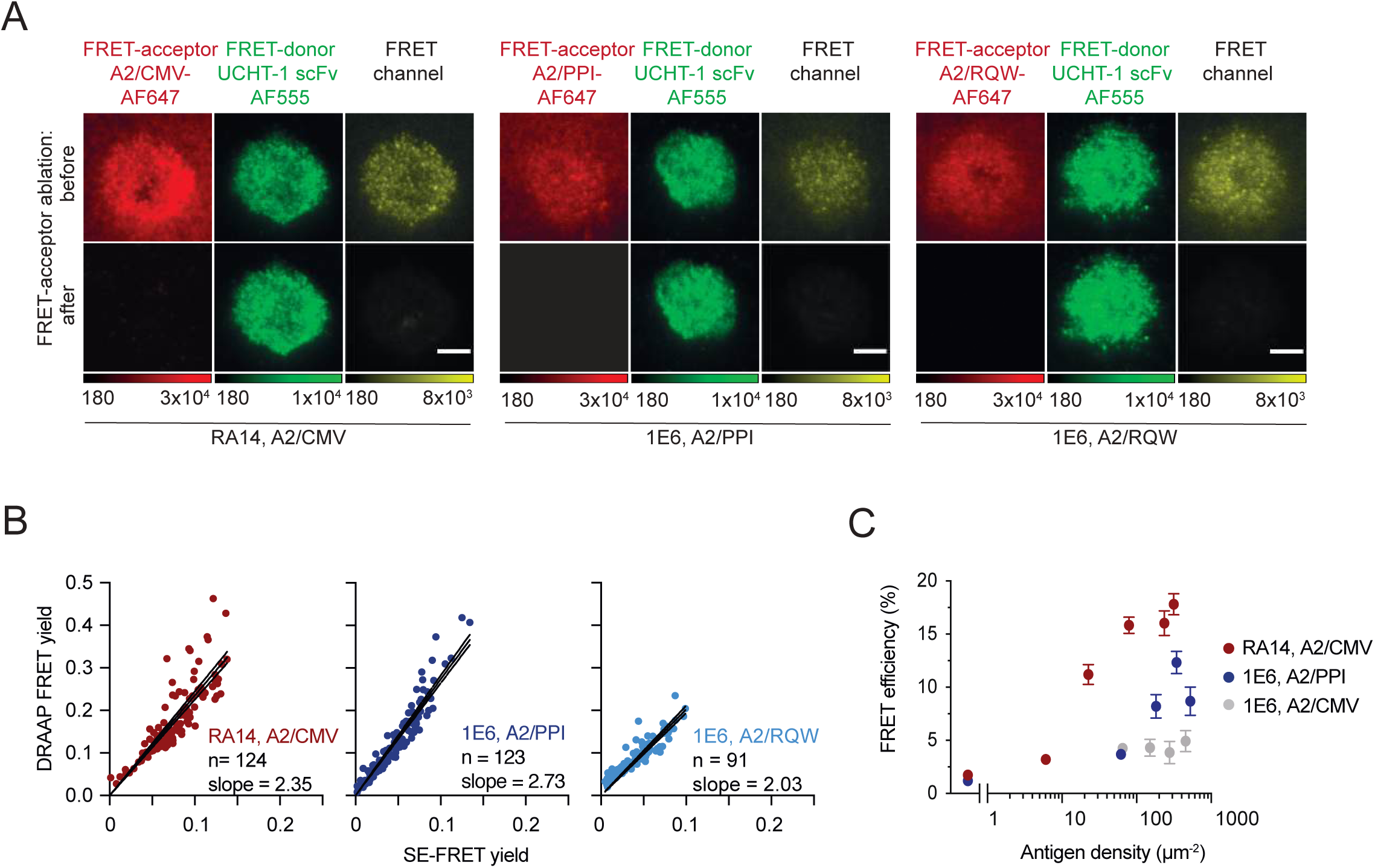
FRET-based system to quantitate TCR-peptide/HLA interactions *in situ*. (**A**) FRET acceptor, FRET donor and FRET channels of RA14 and 1E6 T-cells which have been decorated with UCHT-1 scF_V_-AF555 and which engage SLB-resident peptide/A2 ligands before and after acceptor photobleaching. Images have been acquired in TIRF configuration. Scale bar, 5µm. (**B**) Correlation of FRET yields determined by Fluorescence Sensitized Emission (SE-FRET) and FRET yields measured by FRET Donor Recovery after FRET Acceptor Photobleaching (DRAAP) with the slope of the corresponding linear fit (black line). (**C**) FRET efficiencies as determined by Donor Recovery After Acceptor Photobleaching (DRAAP) in synapses of RA14 and 1E6 T-cells engaging SLB-embedded A2/CMV (red, grey) and A2/PPI (blue) at indicated antigen densities. Data shown are representative of (n=3) independent experiments with the use of T-cells isolated from three different donors.

**Figure S4.**
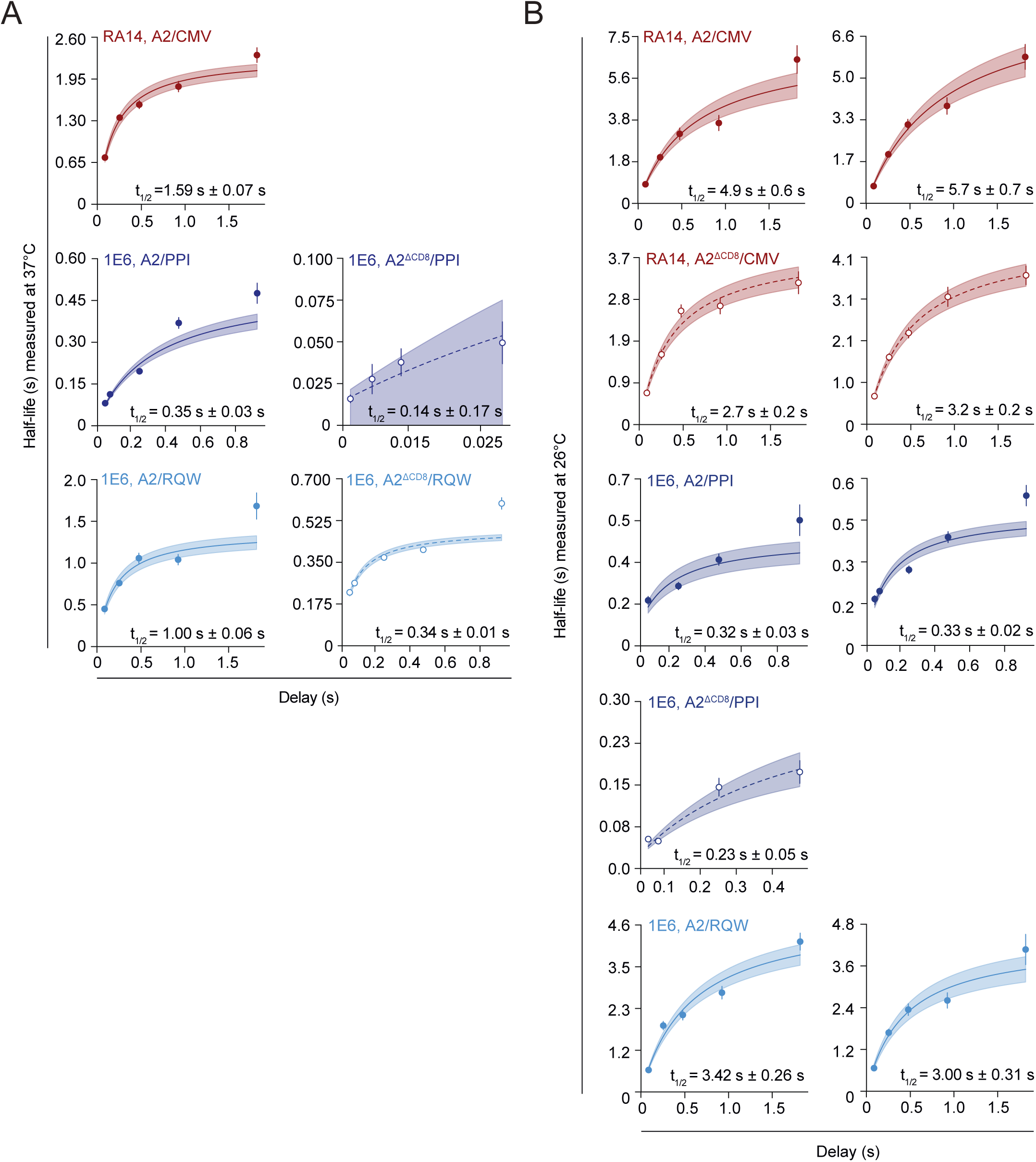
Single molecule FRET measurement of TCR:pMHC binding kinetics at 37°C and 26°C. (**A**) Repetitions of TCR-pMHC lifetime measurements (at 37°C) for RA14 and 1E6 T-cells on SLBs featuring indicated pMHCs. Time delays between recorded images are as indicated (7 ms, 10 ms, 14 ms, 28 ms, 56 ms, 84 ms, 252 ms, 476 ms, 924 ms, 1820 ms). Shaded areas refer to 95% confidence intervals. Measured half-lives and standard deviations (SDev) are indicated. Data are shown as representative of n=2 experiments. (**B**) Non-linear least squares regression fit of measured lifetimes (at 26°C) of synaptic TCR-peptide/HLA interactions recorded for RA14 and 1E6 T-cells scanning SLBs for indicated pMHCs. Time delays between recorded images are as indicated 56 ms, 84 ms, 252 ms, 476 ms, 924 ms and 1820 ms. Shaded areas refer to 95% confidence intervals. Measured half-lives and standard deviations (SDev) are indicated. Data are shown as representative of n=2 experiments.

